# Epinephrine inhibits PI3K alpha via the Hippo kinases

**DOI:** 10.1101/2022.07.19.500601

**Authors:** Ting-Yu Lin, Shakti Ramsamooj, Katarina Liberatore, Louise Lantier, Neil Vasan, Kannan Karukurichi, Seo-Kyoung Hwang, Edward A. Kesicki, Edward R. Kastenhuber, Thorsten Wiederhold, Tomer M. Yaron, Mengmeng Zhu, Yilun Ma, Marcia N. Paddock, Guoan Zhang, Benjamin D. Hopkins, Owen McGuinness, Robert E. Schwartz, Lewis C. Cantley, Jared L. Johnson, Marcus D. Goncalves

## Abstract

The phosphoinositide 3-kinase, p110α, is an essential mediator of insulin signaling and glucose homeostasis. We systematically interrogated the human serine, threonine, and tyrosine kinome to search for novel regulators of p110α and found that the Hippo kinases phosphorylate and completely inhibit its activity. This inhibitory state corresponds to a conformational change of a membrane binding domain on p110α, which impairs its ability to engage membranes. In human primary hepatocytes, cancer cell lines, and rodent tissues, activation of the Hippo kinases, MST1/2, using forskolin or epinephrine is associated with phosphorylation and inhibition of p110α, impairment of downstream insulin signaling, and suppression of glycolysis and glycogen synthesis. These changes are abrogated when MST1/2 are genetically deleted or inhibited with small molecules. Our study reveals a novel inhibitory pathway of PI3K signaling and a previously unappreciated link between epinephrine and insulin signaling.

## INTRODUCTION

The binding of insulin to the insulin receptor initiates a surge of tyrosine phosphorylations that promote membrane recruitment of phosphoinositide 3-kinase (PI3Kα), an intracellular lipid kinase(Chen et al., 2019; Fritsch et al., 2013; Nolte et al., 1996). PI3Kα is composed of a catalytic subunit, p110α, which functions as an obligate heterodimer with the regulatory subunit, p85α. Binding of the regulatory subunit to phosphotyrosines relieves its inhibitory contacts on p110α and localizes the complex to the plasma membrane(Fruman et al., 2017), where p110α then engages the lipid bilayer and phosphorylates the D3-positions of the resident phosphoinositide lipids to set off signaling cascades that guide the cellular actions of insulin, including glucose uptake, enhanced glycolysis, and glycogen synthesis(Hopkins et al., 2020; Manning and Toker, 2017).

The glycemic actions of insulin and PI3Kα are opposed by counter-regulatory hormones like glucagon and epinephrine (Epi). Epi is released from the adrenal gland in response to stresses like hypoglycemia. It then binds to β-adrenergic receptors (β_2_-AR) in the liver to stimulate glycogenolysis and impair the actions of insulin(Cori et al., 1930; Deibert and DeFronzo, 1980; Dufour et al., 2009). The molecular pathways by which Epi acutely blocks insulin action are unclear(Battram et al., 2007; Cori and Cori, 1929; Deibert and DeFronzo, 1980; Dufour et al., 2009; Rizza et al., 1980).

Most of what is known about the regulation of p110α has come from experimental work with accessible cellular model systems, using indirect readouts as a surrogate for PI3Kα activation. We have taken a more direct approach and asked if the production of phosphatidylinositol 3,4,5-triphosphate (PIP_3_), the product of p110α, can be regulated through phosphorylation by a broad sample of the human protein kinome. This approach has unveiled a novel inhibitory pathway of PI3K signaling that is used by Epi to block insulin signaling.

## RESULTS

### The Hippo kinases inhibit p110α through phosphorylation of threonine 1061 at its C-terminus

To investigate phospho-regulation of PI3Kα (i.e., the p110α/p85α complex), we developed an *in vitro* reconstitution assay to screen for the effects of recombinant protein kinases on its catalytic activity. Recombinant, full length p110α was purified from human suspension cultures as a complex with polyhistidine-tagged p85α(Dickson et al., 2013; Sun et al., 2011; Vasan et al., 2019). This complex was then pre-incubated with a panel of 31 functionally diverse human protein kinases from the major branches of the protein kinase evolutionary tree(Manning et al., 2002), and we evaluated the ability of PI3Kα to phosphorylate phosphatidylinositol 4,5-bisphosphate (PIP_2_), its endogenous substrate (Figs. 1A/B). Of the 31 kinases, only MST1 (encoded by *Stk4*) had a major effect on PI3Kα catalytic activity, causing nearly complete inhibition with dose-dependence at the level of the protein kinase (Fig. 1C).

**Fig. 1.**
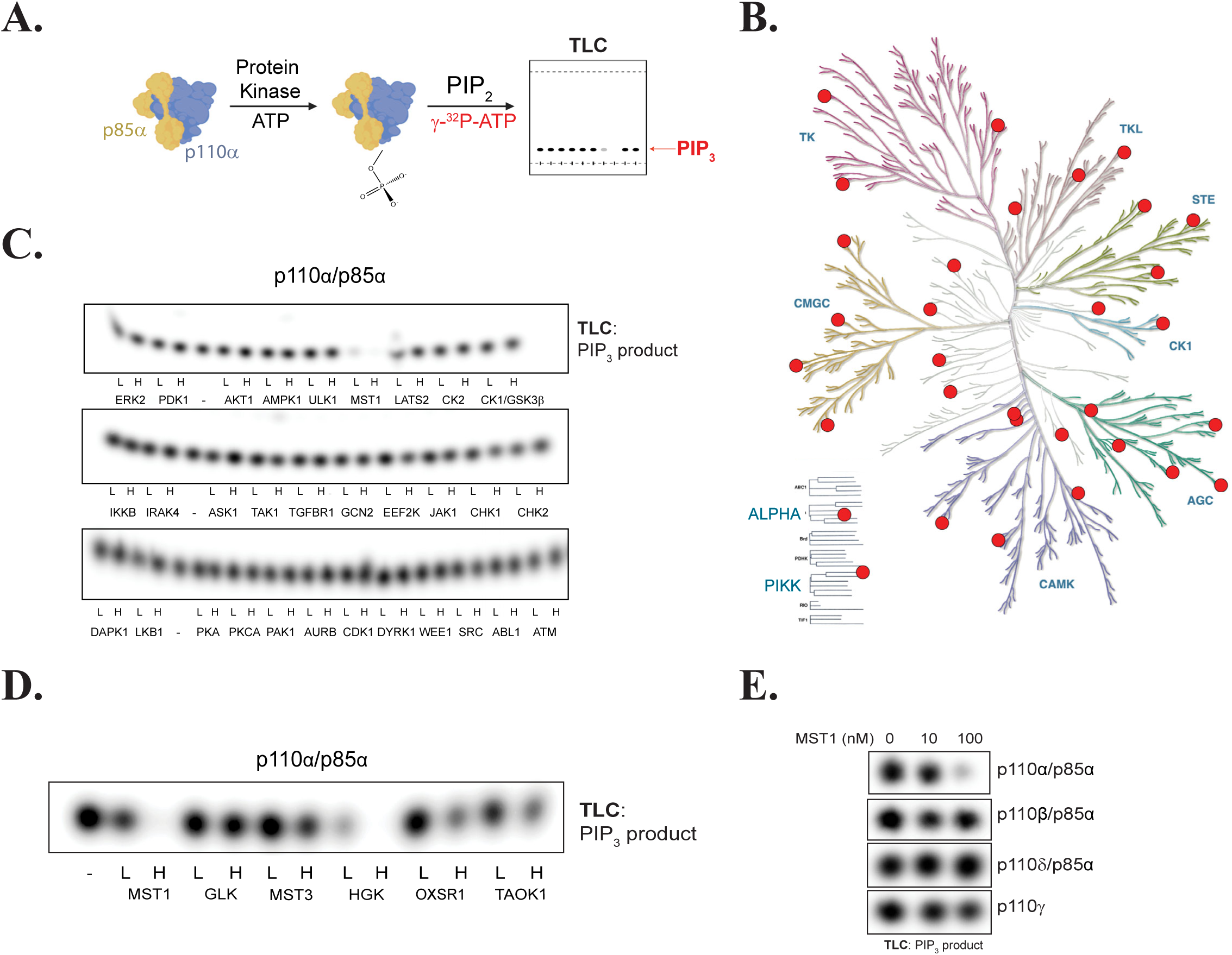
Kinase screen identifies GCK kinases as negative regulators of PI3K*α* activity. (A) Experimental schema. (B) Dendrogram of the human protein kinome that highlights the kinases selected for our screen. (C) Thin layer chromatography (TLC)-autoradiography of [^32^P]PIP produced by recombinant PI3Kα complexes (p110α/p85α, 500 nM) after incubation with the indicated recombinant protein kinases at 10 nM (L) or 100 nM (H) concentrations or with enzyme storage buffer (-). (D) PIP_2_ kinase assays with PI3Kα after incubation with subset of the GCK family, performed as described in (C). (E) PIP_2_ kinase assays with the class I PI3K isoforms (PI3Kα, PI3Kβ, PI3Kγ, and PI3Kδ, 500 nM) after incubation with MST1.

MST1, the human ortholog of hippo (hpo) in *Drosophila*, is a growth suppressing kinase in the group II germinal center kinase (GCK II) family, and a core member of the well-conserved Hippo pathway(Dong et al., 2007; Harvey et al., 2003; Wu et al., 2003). Because the GCK family members have overlapping cellular functions and similar substrate specificities(Meng et al., 2015; Miller et al., 2019), we tested whether additional GCK family members (Fig. S1) can inhibit PI3Kα’s catalytic activity. We found that HGK, OXSR1, TAOK1, and MST3 could also inhibit PI3Kα, suggesting that PI3Kα may be under broader regulation by the GCK family (Fig. 1D). Next, we investigated MST1’s ability to inhibit other class I PI3K isoforms. MST1 had little effect on the catalytic activities of p110β, p110δ, and p110γ, indicating a selective regulation of p110α (Fig. 1E). Like the p110α preparation, the p110β and p110δ proteins were prepared as complexes with p85α. Given their insensitivity to MST1 inhibition, we inferred that MST1 was directly phosphorylating p110α, and not p85α.

To locate the MST1 phosphorylation site(s) on p110α, we profiled MST1’s *in vitro* substrate specificity using positional scanning peptide arrays (PSPA). This technique utilizes a combinatorial peptide library that systematically substitutes each of 22 amino acids (20 natural amino acids plus phospho-Thr and phospho-Tyr) at nine positions surrounding a central phospho-acceptor containing equivalent amounts of Ser and Thr (Hutti et al., 2004). MST1 preferentially phosphorylated threonine over serine and favored aliphatic amino acids at position +1, aromatic amino acids at position -2, and positively charged amino acids at position +2 (Figs. 2A). The PSPA data were converted into a positional specific scoring matrix (PSSM) and used to score all threonine residues in the amino acid sequence of p110α. The most favorable *in silico* MST1 threonine target was T1061, located in p110α’s C-terminal tail (C-tail), a disordered region spanning residues 1049 to 1068 (Fig. 2B). To validate this finding, we incubated recombinant PI3Kα with MST1 and analyzed peptide fragments by mass spectrometry. This approach confirmed that phosphorylation of T1061 increased after MST1 treatment (Fig. S2A). Furthermore, we generated the recombinant threonine-substituted phospho-null mutant, T1061A, of p110α and found its catalytic activity to be completely unaffected by MST1 and HGK (Fig. 2C and S2B).

**Fig. 2.**
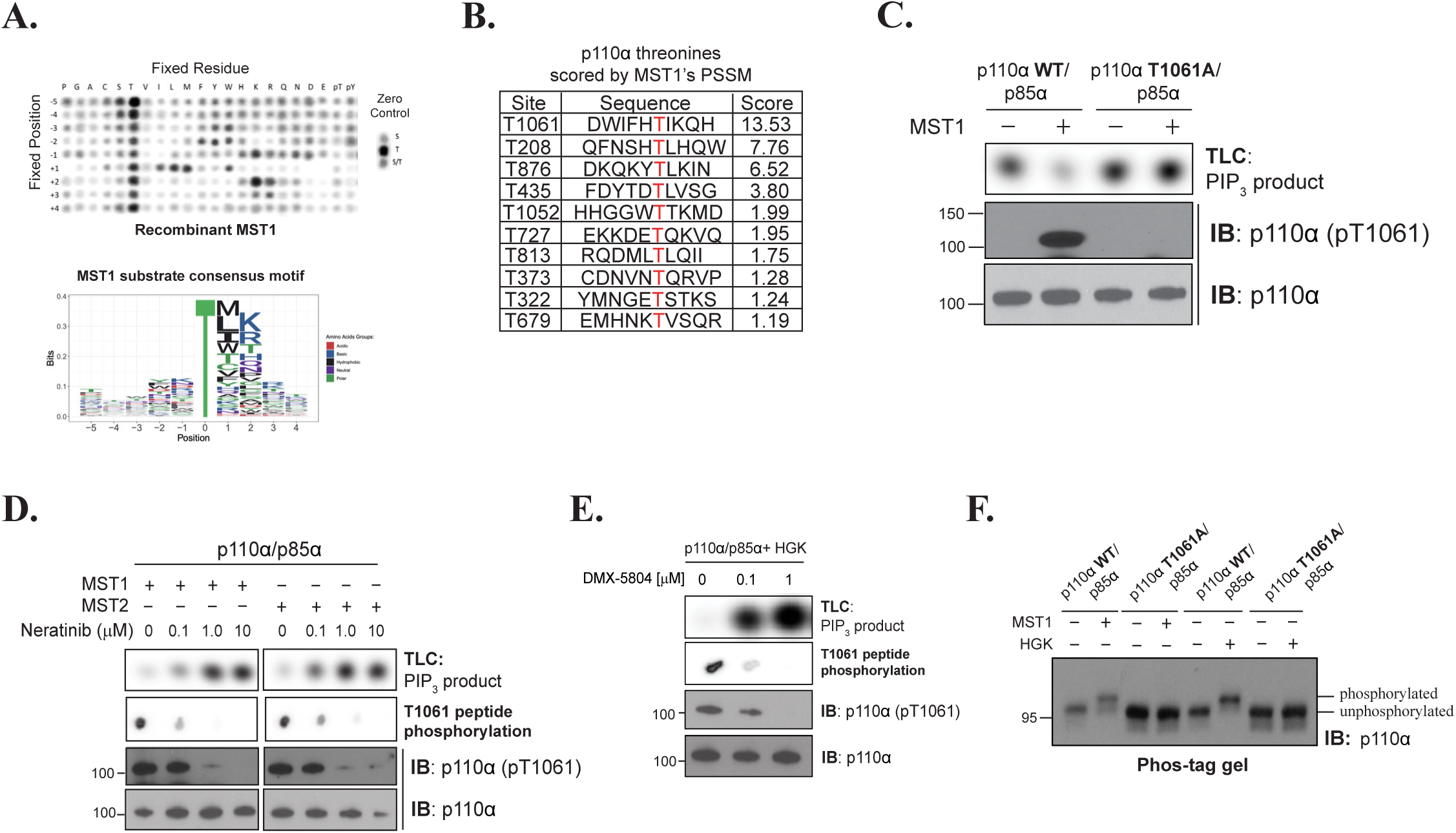
MST1/2 and HGK inhibit catalytic activity of p110*α* through phosphorylation at T1061. (A) (Top) Peptide phosphorylation by MST1 to characterize its substrate consensus motif. Positional scanning peptide arrays were utilized, where 22 residues (20 amino acids + 2 PTM residues) were scanned across nine neighboring positions of the phospho-acceptor. The zero controls (*righthand side*), consisting of serine only, threonine only, or a 1:1 mixture of both were examined as phospho-acceptors. Phosphorylation was measured by autoradiography. (Bottom) Sequence logo of the substrate consensus motif of MST1 as determined in top panel. Letter height is proportional to favorability of corresponding amino acid. (B) p110α’s threonine residues scored by MST1’s PSSM obtained from (A). (C) Incubation of p110α (WT)/p85α and p110α (T1061A)/p85α with MST1. Top: Autoradiography of [^32^P]PIP production by p110α. Bottom: Immunoblots of total p110α and pT1061 p110α. (D) Incubation of MST1 or MST2 with increasing concentrations of Neratinib, followed by incubation with PI3Kα. Top: Autoradiography of [^32^P]PIP production by p110α. Second from top: Autoradiography of T1061-modeled peptidesubstrate peptide phosphorylation by MST1 or MST2. Bottom: Immunoblots of total p110α and pT1061. (E) Repeat of (D) using HGK and and its specific inhibitor DMX-5804. (F) Immunoblot of p110α (WT and T1061A) on Phos-tag gel after treatment with MST1 or HGK.

Next, we generated and validated polyclonal antibodies against a p110α peptide containing phosphothreonine at 1061 (pT1061). The antibodies detected wild-type p110α only after MST1-treatment and failed to identify MST1-treated T1061A, confirming their utility for readout of pT1061 (Fig. 2D). We also showed that blocking MST1/2 or HGK activity using Neratinib or DMX-5804, respectively, reversed the inhibition of PI3Kα activity and reduced pT1061 phosphorylation (Fig. 2D/E). Finally, we performed SDS-PAGE using Phos-tag gels to measure the extent of phosphorylation of p110α by MST1 and HGK(Kinoshita et al., 2006). Following exposure to MST1 or HGK, the protein band blotted by an anti-p110α antibody exhibited a delay in migration, suggesting stoichiometric modification of p110α (Fig. 2F). In contrast, the T1061A protein migrated alongside that of unphosphorylated wild type p110α in both control and MST1/HGK-treated samples, indicating that pT1061 is the prominent phosphorylation event. Together, these data indicate that PI3K catalytic activity is inhibited through phosphorylation of T1061 on p110α by the Hippo kinases.

### Phosphorylation of T1061 inhibits p110α’s interaction with membranes

To investigate the stability of the PI3K complex following inhibition at pT1061, we performed thermal shift assays. Recombinant PI3Kα complexes were phosphorylated by MST1 and subjected to increasing temperatures. Denatured proteins were removed by centrifugation and the remaining soluble proteins were examined by Western blotting. MST1-mediated phosphorylation led to a significant stabilization of wild type p110α, and not p110α (T1061A), at higher temperatures (Fig. 3A), indicating that pT1061 induces a stabilizing conformational change. The stabilizing action of pT1061 suggests an ordering of the C-tail. Only ∼15% (8/55) of publicly available structures of PI3Kα can resolve T1061 and when resolved its side chain faces the solvent(Barsanti et al., 2015; Furet et al., 2013; Han et al., 2016; Heffron et al., 2016; Hon et al., 2012; Mandelker et al., 2009). In general, the tail adopts two conformations: one that is directed towards the catalytic site in a ‘frontside conformation,’ or one where T1061 undergoes a ∼42 angstrom shift to the opposite side of the protein in a ‘backside conformation’ (Figs. 3B/C)(Walker et al., 2000; Walker et al., 1999). We generated and purified a phosphomimetic mutant of p110α (T1061E) and showed that it partially inhibits PI3K activity (Fig S3 A-C). To determine the structural basis for PI3Kα inhibition by T1061 phosphorylation, we solved the crystal structure of the p110α T1061E protein in complex with the nSH2-iSH2 domains of p85α and catalytic inhibitor GDC-0077, at 2.95 angstrom. The C-tail was resolved up to and not beyond the phosphomimetic glutamate and adopted the backside conformation observed in previous structures of wild type p110α (Figs. 3B/C). However, the side chain of E1061, unlike T1061 from previous structures, did not face the solvent but was directed into the protein. The carboxylate of E1061 was positioned to form an ion-dipole interaction with the amino terminal end of the helical dipole generated by helix kα11, encompassing residues 1031-1048(Hol et al., 1978). These structural findings suggest an allosteric model of inhibition by T1061 phosphorylation.

**Fig. 3.**
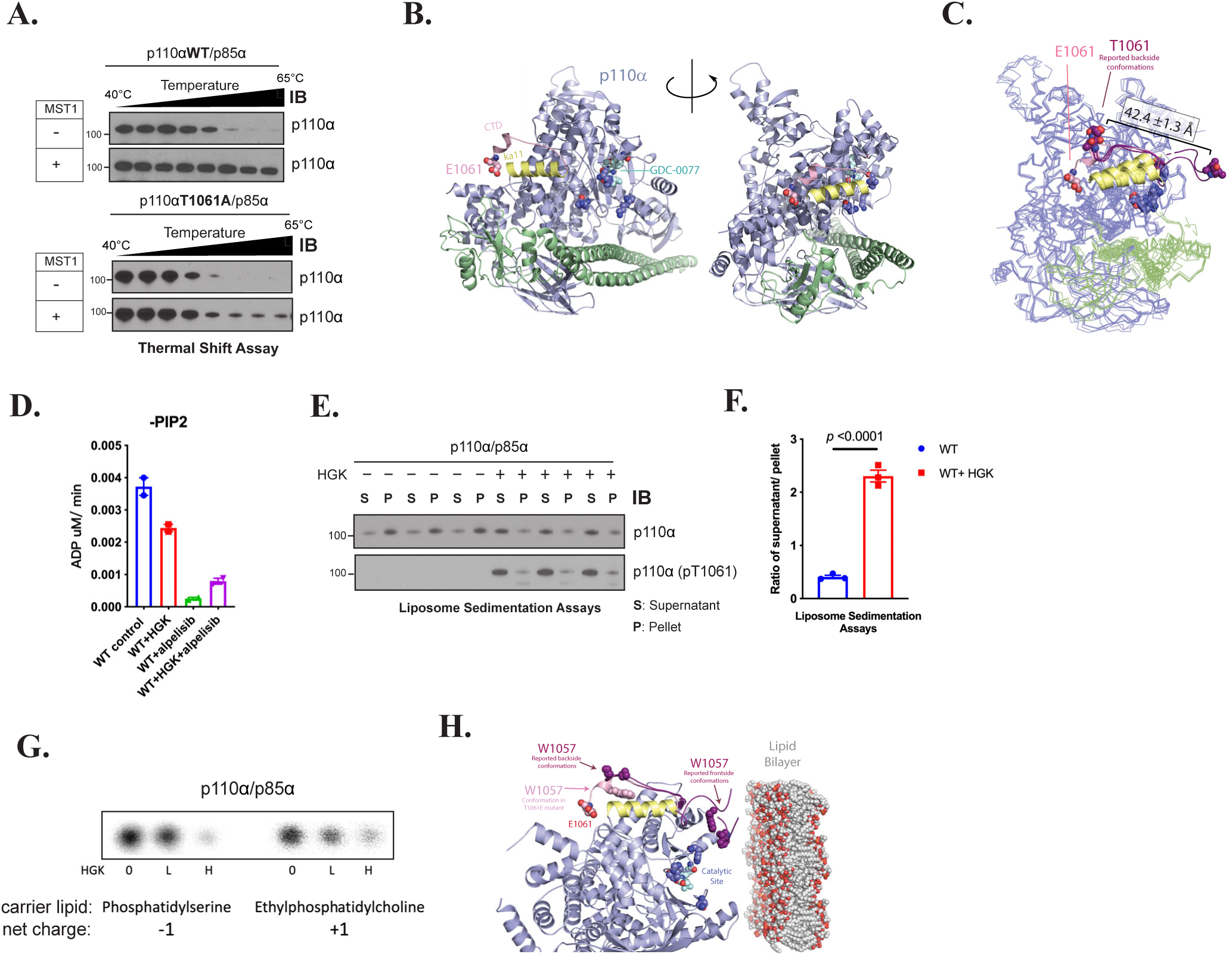
Phosphorylation at T1061 prevents p110*α*’s association with membranes. (A) Thermal shift assays of p110α (WT and T1061A) after MST1 treatment. (B) Structure of p110α (T1061E)/p85α-niSH2. Residues 1049-1061, constituting the C-tail, are displayed in pink. E1061’s side chain is displayed as spheres. Residues 1032-1048, constituting helix ka11, are displayed in yellow. Catalytic residue side chains K776, H917, and H936^35^ are displayed as dark blue spheres and the bound catalytic inhibitor GDC-0077 is displayed as teal spheres. (C) Overlay of the crystal structure of p110α (E1061) with six reported p110α (WT) structures where threonine 1061 was resolved. C-tails from frontside conformations (pdb ids: 4a55 and 5dxh) and backside conformations (pdb ids: 4jps, 4waf, 4zop, and 5fi4) of reported structures are shown in purple. (D) ADP Glo measurements of ATPase activity of PI3Kα -/+ HGK and -/+ 1 μM alpelisib. (E) Liposome sedimentation assays of PI3Kα -/+ HGK, shown as immunoblots of p110α or p110α (pT1061) recovered from membrane enriched pellet (P) and supernatant (S). (F) Quantification of densitometries from (E), as ratios of p110α or pT1061 recovery from supernatant over pellet. Data were represented as means ± SEMs. Significance was calculated using student’s t test (N=3). (G) Activity assays of PI3Kα on PI in anionic and cationic liposomes after incubation with HGK. [^32^P]PIP_3_ products were resolved by TLC and measured by autoradiography. (H) Overlay of p110α (T1061E)’s crystal structure (C-tail in pink) with the 6 reported p110α(WT) structures (in purple, C-tails shown only) selected in Figure 2C. The coloring scheme corresponds to Figure 2C. W1057 sidechains are represented as spheres. Catalytic site is indicated by residues K776, H917, and H936 (dark blue) and bound GDC-0077 (teal), shown as spheres. The phospholipid membrane model was obtained from RCSB PDB (pdb id: 2mlr).

Our crystal structure indicates that phosphorylation of p110α T1061 diverted its C-tail away from the catalytic site, so we investigated how this change causes inhibition. Given that p110α is activated chiefly through enhancement of its catalytic turnover rate and/or membrane binding(Burke, 2018; Carson et al., 2008; Chaussade et al., 2009; Gkeka et al., 2014; Gymnopoulos et al., 2007; Huang et al., 2007; Mandelker et al., 2009; Miled et al., 2007; Shekar et al., 2005; Vasan et al., 2019), we considered both of these possibilities for MST1-mediated phosphorylation. In the absence of substrate, p110α exhibits a basal ATPase activity that is a small fraction of the rate of phosphate transfer to PIP_2(Hon et al., 2012)_. This basal ATPase activity was inhibited by the ATP competitive PI3Kα-specific inhibitor, alpelisib, indicating that it is intrinsic to p110α(Furet et al., 2013). Phosphorylation of p110α at T1061 caused a modest reduction in this activity, indicating that the phosphorylation at T1061 does not significantly affect ATP binding and phosphate transfer to water (Fig. 3D).

Next, we determined if p110α phosphorylation affects its interaction with membranes using liposomes that were modeled after the inner leaflet of the plasma membrane. The liposomes were incubated with p110α and pelleted by centrifugation. The amounts of p110α recovered from supernatant and pellet were then quantified. Under control conditions, p110α was mostly lipid-bound (Fig. 3E). However, in HGK-treated samples, we saw an appreciable shift into the supernatant, measured as a 6-fold increase in the ratio of p110α recovered from supernatant versus pellet (Figs. 3E/F). These data indicate that T1061 phosphorylation impairs membrane binding. Indeed, MST1 treatment and introduction of glutamate to position 1061 also reduced membrane binding of p110α (Figs. S4A/B). Given that the C-tail interacts with membranes, we examined the possibility that membrane dissociation is driven by electrostatic repulsion between the phosphorylated C-tail and the negatively charged membrane surface. To test this, cationic liposomes containing phosphatidylinositol (PI) were prepared as substrates. HGK-treatment reduced p110α‘s conversion of PI to PI3P in both our anionic control liposomes and the cationic liposomes, indicating that charge repulsion is not a significant contributor (Fig. 3G). The C-tail of p110α contains a hydrophobic patch, composed of residues 1057-1059, that has been reported to be essential for membrane binding(Hon et al., 2012). Within this patch, tryptophan 1057 is universally conserved across all examined p110α orthologs (Fig. S4C). In the reported structures of wild type p110α, the W1057 sidechain only resolves in the frontside conformations of the tail to form part of the predicted membrane binding interface. Our structure of the T1061E phosphomimetic mutant is a novel case where W1057 can be stabilized and resolved in the backside conformation where it cannot directly contribute to membrane binding (Fig 3H). Our findings are consistent with a model where T1061 phosphorylation acts by redirecting the C-terminus away from the catalytic face of p110α, sequestering a membrane binding domain, and thereby reducing its ability to engage membranes and phosphorylate PIP_2_.

### Activation of adenylyl cyclase promotes phosphorylation and inhibition of p110α

We proceeded to explore the biological implications of p110α regulation by the Hippo kinases. Recent work has identified the Hippo pathway as a downstream branch of G-protein-coupled receptor (GPCR) signaling.(Yu et al., 2012) Stimulation of the G_αs_-coupled receptors or adenylyl cyclase using receptor agonists or forskolin (FSK), respectively, results in activation of the PKA and Hippo kinase signaling pathways(Yu et al., 2013). To determine if p110α can be regulated by the Hippo pathway under these contexts, we first treated primary mouse hepatocytes with FSK(Yu et al., 2012). FSK strongly induced phosphorylation of p110α at T1061 as well as PKA activity as denoted by the increase of CREB phosphorylation at S133 (Fig. 4A). We confirmed this finding in multiple cell lines (Figs. S5A-C). To further clarify the effects of pT1061 on PI3K activity, we treated freshly isolated primary human hepatocytes with insulin followed by FSK. Insulin robustly stimulated p110α activity as determined by AKT phosphorylation at T308 and S473, and the addition of FSK led to increased PKA activity, p110α phosphorylation at T1061, and a reduction of AKT phosphorylation (Fig. 4B). This was also demonstrated in MCF10A and AML12 cells (Figs. S5D-E). The activation of PI3K results in multiple downstream feedback pathways so we directly examined the catalytic activity of p110α using an *in vitro* activity assay with immunopurified enzyme taken from cells treated with insulin and FSK (Figs. 4C/D). In the presence of insulin stimulation, the production of PIP_3_ was increased over 100-fold, and this was greatly reduced when FSK was added. The phosphorylation and inhibition of p110α by FSK in HEK293A cells also correlated with suppression of the extracellular acidification rate (ECAR), suggesting acute reductions in glycolysis (Figs. 4E/F). Taken together, these results demonstrate that FSK-induced adenylyl cyclase activity leads to phosphorylation and inhibition of p110α and glucose metabolism.

**Fig. 4.**
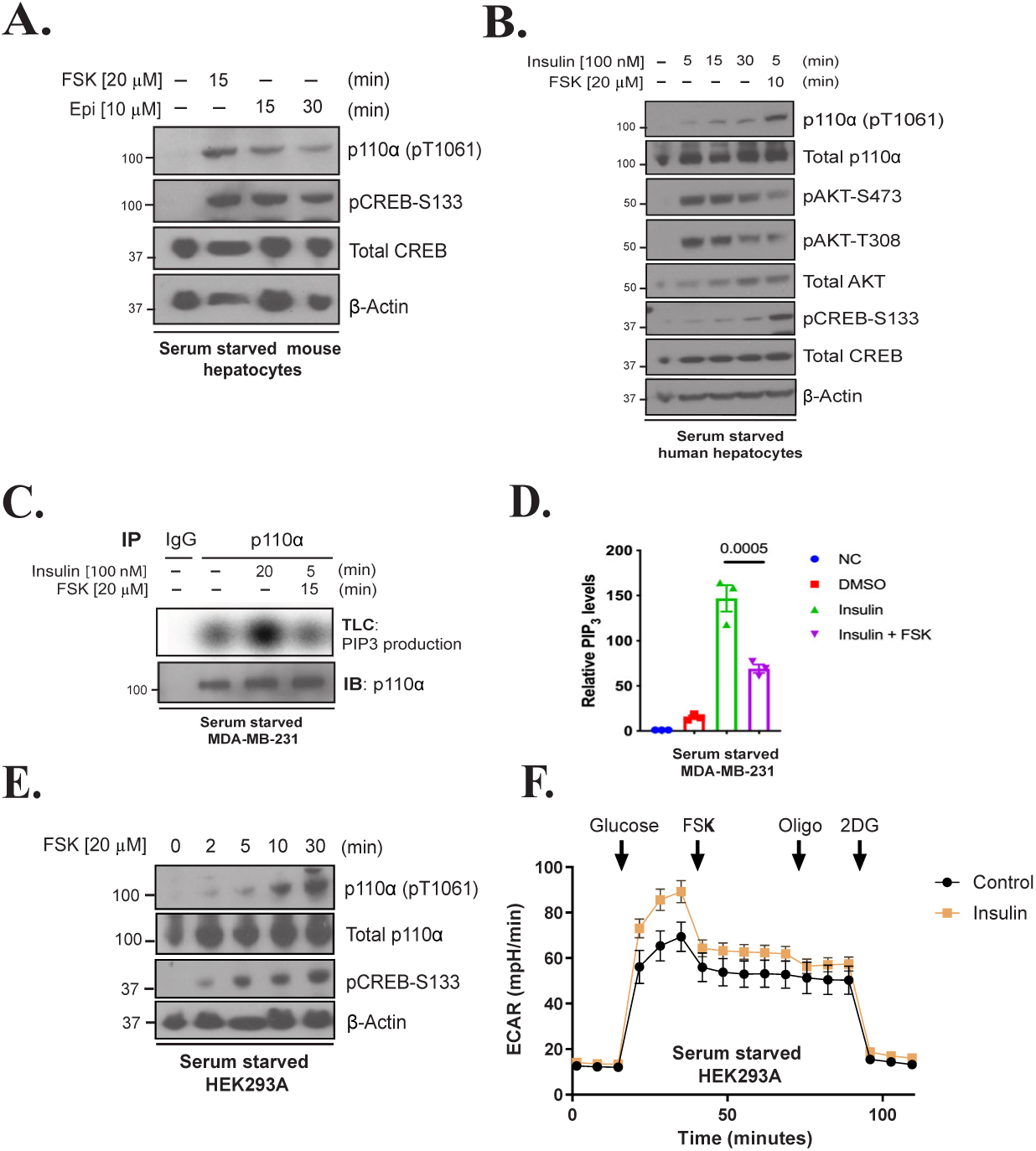
Forskolin and epineprine exposure phosphorylate and inhibit p110*α* in cells. (A) Immunoblot for the indicated proteins using lysates from primary mouse hepatocytes that were serum starved 12 h before stimulation with forskokin (FSK, 20 μM) for 15 min or epinephrine (Epi, 10 μM) for 15 and 30 min. (B) Immunoblot for the indicated proteins using lysates from primary human hepatocytes that were serum starved for 12 h, and treated with insulin [100 nM] for different time points (0, 5, 15, 30 min) or insulin followed by FSK [20 μM] for 10 min. (C) Top: Radioautograph of a TLC separation demonstrating PIP_3_ production of endogenous p110α that was immunoprecipitated from serum starved MDA-MB-231 cells treated with vehicle, insulin [100 nM] or insulin plus FSK [20 μM] for 15 min. Bottom: Corresponding immunoblot for p110α using the same immunoprecipitate lysate. (D) Quantification of the radioautograph from (C) averaged over three independent experiments. Means ± SEM. Comparisons made using ANOVA with Tukey’s multiple comparisons post-test (N=3). \*\*\**p*< 0.001. (E) Immunoblot for the indicated proteins using lysates from HEK293A cells that were serum starved for 2 h before stimulated with FSK [20 μM] for 0, 2, 5, 10, and 30 min. (F) The extracellular acidification rate (ECAR) was monitored in serum starved HEK293A cells with or without insulin [0.1 μM] pre-treatment for 1 h. Arrows indicate injection of glucose [10 mM], FSK [20 μM], oligomycin [Oligo, 1 μM], and 2-deoxy-D-glucose [2DG, 50 mM]. N=19

### Epinephrine stimulates p110α phosphorylation at T1061 *in vivo*

Epinephrine (Epi) also stimulates adenylyl cyclase activity and regulates the Hippo pathway through GPCRs including the β_2_-AR, a G_αs_-coupled receptor.(Yu et al., 2013; Yu et al., 2012) Therefore, we hypothesized that Epi could result in phosphorylation and inhibition of p110α. First, we confirmed that Epi induces p110α phosphorylation at T1061 using primary mouse hepatocytes (Fig 4A). Next, we injected wild-type C57BL/6J mice with Epi and collected liver tissue for Western blot analysis. In the Epi-treated mice, we observed a significant increase in the phosphorylation of T1061 in liver (Figs. 5A). This increase was not observed in mice with genetic deletion of the β_1_ and β_2_-AR (Fig. 5B), confirming the importance of β-adrenergic signaling in this response. Next, we exposed mice to several physiologic stressors to induce endogenous Epi release and assess the phosphorylation status of p110α. For example, hypoglycemia is known to induce the sympathoadrenal response and cause an acute and robust release of Epi so we fasted the mice for 18 h. Fasting increased blood Epi levels and significantly increased the abundance of p110α phosphorylation at T1061 in the liver as compared to “Fed” mice that were food restricted for 18 h and then given access to food for 4 h (Figs. 5C-E). The fasted mice also displayed reduced levels of phosphorylated AKT, as expected in conditions of low circulating insulin levels (Fig. S6A). Next, we imposed hyperinsulinemic conditions by treated wild-type mice with insulin until hypoglycemia developed (30 min) and harvested blood and liver tissues. At this time point, Epi levels in the blood were elevated, the abundance of T1061 phosphorylation was significantly increased, and phosphorylation of AKT was reduced, suggesting that Epi blocks PI3K activity in the liver (Figs. 5F-H). Lastly, we fed mice a high-fat diet (HFD), which is known to increase endogenous Epi levels(Gomes et al., 2019). Following 2 weeks of HFD, systemic Epi levels were elevated, and the abundance of pT1061 was increased as compared to mice fed with normal chow (Figs. S6B-D).

**Fig. 5.**
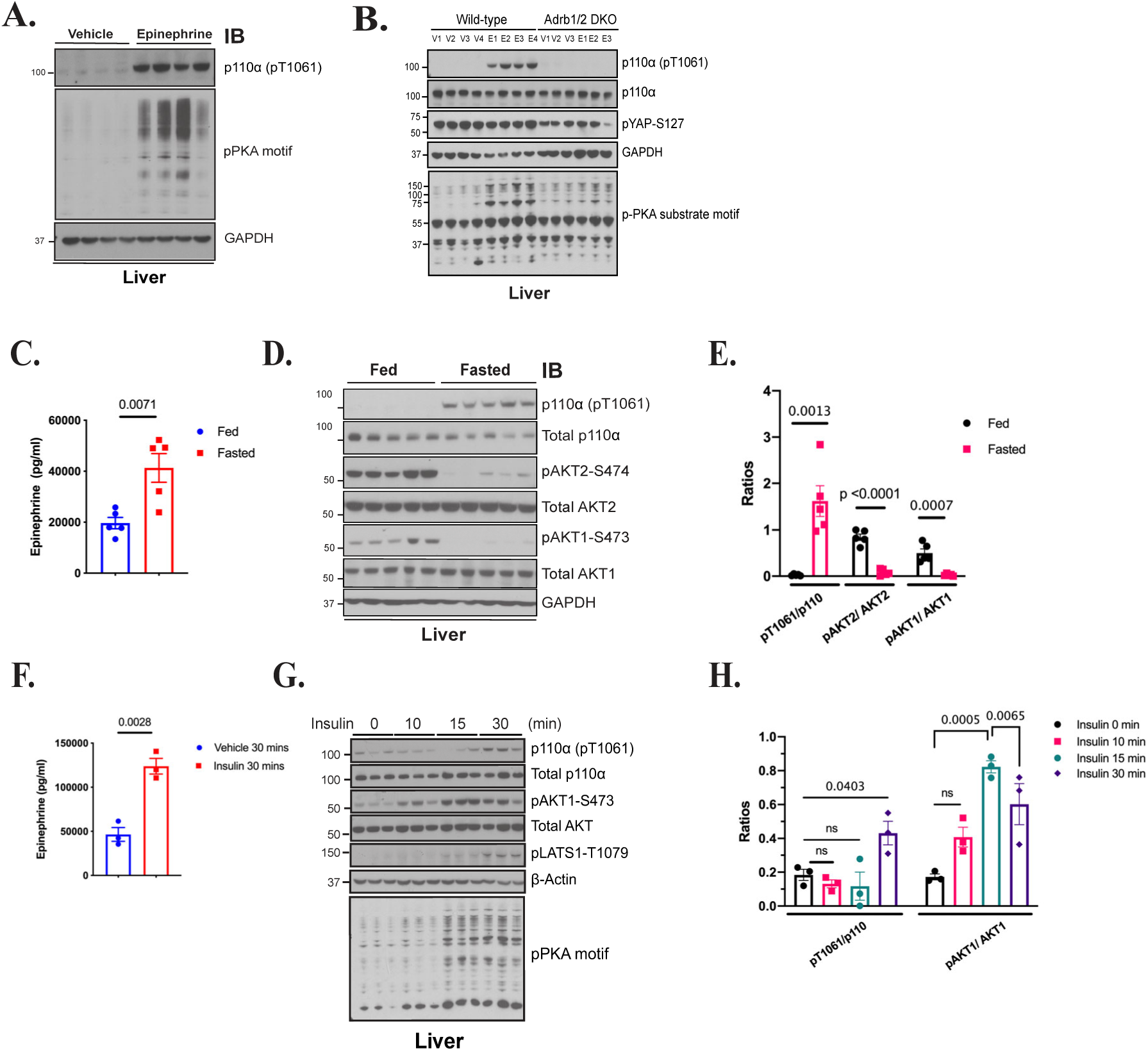
Epinephrine exposure induces phosphorylation and inhibition of p110*α* in the liver. (A) Immunoblot for the indicated proteins using lysates from livers taken from WT mice that were injected with vehicle (normal saline) or epinephrine [0.3 μg/g] for 10 min (n=4). (B) Immunoblot for the indicated proteins using lysates from livers taken from WT and β1/β2 double knockout mice (Adrb1/2 DKO) treated with vehicle (V, saline) or epinephrine (E, Epi) via I.P. injection for 10 min. (C) Plasma Epi levels in WT mice that were fasted for 18 h (Fasted) and then euthanized or provided food for 4 h (Fed). N=5. (D) Immunoblot for the indicated proteins using lysates from liver taken from WT Fed and Fasted mice (n=5). (E) Quantification of the ratios of pT1061 to total p110α, pAKT2 to total AKT2 and pAKT1 to total AKT1 using band intensity from (D). N=5. (F) Plasma Epi levels in WT mice that were injected with insulin [0.75 mIU/g] for 0 and 30 min. N=3. (G) Immunoblot for the indicated proteins using lysates from livers taken from WT mice injected with insulin [0.75 mIU/g] for different time periods (0, 10, 15 and 30 min). (H) Quantification of the ratios of phosphorylated p110α (pT1061) to total p110α and pAKT1 to total AKT1 using band intensity from (G). N=3. Lines and bars indicate Mean ± SEM. For C, E, F comparisons made via student’s t-test. For H, comparisons made via ANOVA with Dunnett’s post-test comparing to insulin time 0 min.

In addition to the effects on the liver, Epi increases rates of adipocyte lipolysis leading to the release of non-esterified fatty acids (NEFA) and glycerol, which can then be used by tissues such as the liver. To determine if the lipolytic products from adipose tissue contribute to p110α phosphorylation, we injected Epi into mice with adipocyte-specific deletion of adipose triglyceride lipase (ATGL^aKO^), an enzyme that catalyzes the hydrolysis of triacylglycerols to diacylglycerols. Epi appropriately activated PKA signaling in the adipose tissue of ATGL^aKO^, however there was no rise in NEFA or glycerol in the blood (Fig. S7A-D). Nevertheless, Epi increased the abundance of pT1061 in the liver suggesting that the lipolytic products to do not contribute to this inhibitory signaling event (Fig. S7E).

In order to more stringently assess the effects of Epi on glucose homeostasis, rats were infused with insulin and glucose in order to maintain euglycemic conditions for 90 min prior to the introduction of either vehicle (saline) or Epi (0.75 µg/ kg/ min) for 10 or 30 minutes. At 30 minutes, Epi infusion reduced the glucose infusion rate (GIR), increased pT1061 in hepatic lysates, and increased blood Epi levels (Fig. S8). This increase was associated with markers of MST1/2 activation including LATS1 phosphorylation, and a reduction in markers of PI3K signaling (Fig. S8B/C). Taken together, these experiments demonstrate that endogenous or exogenous Epi stimulation results in MST1/2 activation, which is concomitant with p110α phosphorylation at T1061 and reduced PI3K activity.

### The Hippo kinases, MST1/2, regulate p110α phosphorylation in cells and tissues via PKA

Next, we examined if the Hippo kinases play a role in the phosphorylation of p110α. Primary mouse hepatocytes were treated with FSK in the absence or presence of a selective MST1/2 inhibitor, XMU-MP-1(Fan et al., 2016). This agent significantly reduced FSK and Epi-induced phosphorylation of T1061 (Figs. 6A and S9A). The Epi-induced increase in pT1061 was also significantly reduced when the hepatocytes were pretreated with a non-specific β-adrenergic inhibitor (Propranolol HCl), and the PKA inhibitor, H89 (Fig. S9A). The dependance on PKA was confirmed using another PKA inhibitor, AT13148, and by overexpressing a natural PKA inhibitor (PKIA) in HEK293 cells (Figs. S9B-C). These results implicate PKA as a critical node connecting Epi and MST1/2 to PI3K.

**Fig. 6.**
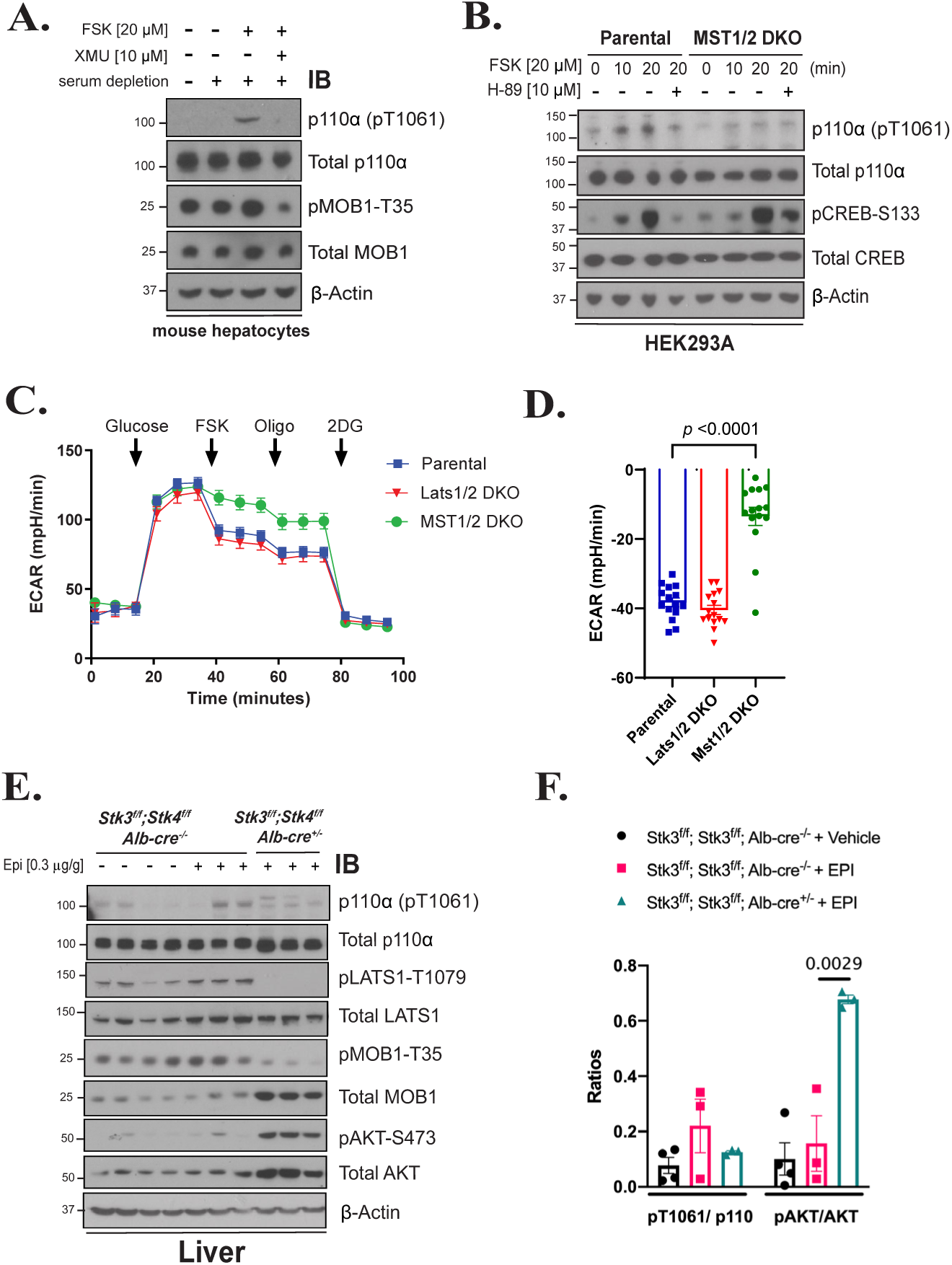
Loss of MST1/2 activity in cells and liver reduces p110*α* phosphorylation. (A) Immunoblot for the indicated proteins using lysates from serum starved primary mouse hepatocytes isolated from WT mice that were stimulated with vehicle (DMSO), FSK [20 μM], or FSK and the MST1/2 inhibitor, XMU [10 μM]. (B) Immunoblot for the indicated proteins using lysates from HEK293A parental cells or a line with CRISPR deletion of MST1 and MST2 (MST1/2 DKO) that were serum starved for 2 h, and pretreated with vehicle (DMSO) or PKA inhibitor, H-89 [10 μM] for 1 h before stimulating with FSK [20 μM] for 10 or 20 min. (C) The extracellular acidification rate (ECAR) was monitored in HEK293A Parental, MST1/2 DKO and Last1DKO cells for 1 h. Arrows indicate injection of glucose [10 mM], FSK [20 μM], oligomycin [1 μM], and 2-deoxy-D-glucose [50 mM]. (D) The suppression (%) of ECAR following the addition of FSK [20 μM] using data from C. N= 15 (E) Immunoblot for the indicated proteins using lysates from livers taken from animals (Stk3^f/f^,Stk4^f/f^, Alb-Cre ^-/-^ and Stk3^f/f^; Stk4^f/f^, Alb-Cre^+/-^) that were injected with vehicle (normal saline) or Epi [0.3 μg/g]. (F) Quantification of the ratios of pT1061 to total p110α, and pAKT to total AKT using band intensity from (E). In C, D, and F, data is represented as Mean ± SEMs. Comparisons made via two-way ANOVA with Sidak’s multiple comparisons post-test (D) and ANOVA with Turkey’s multiple comparisons post-test (F).

We complemented the pharmacologic inhibition of MST1/2 with experiments using cells with CRISPR deletion of MST1/2 (MST1/2 DKO)(Meng et al., 2015). Following treatment with FSK, the MST1/2 DKO and parental cells displayed increased PKA signaling as determined by the phosphorylation of CREB, and an increase in phosphorylation of T1061; however, the phosphorylation T1061 was greatly reduced in MST1/2 DKO cells (Fig. 6B). We next determined if these changes in signaling correspond to changes in PI3K-mediated glucose metabolism using ECAR as a readout. Both MST1/2 DKO cells and cells deficient in Lats1/2 (Lats1/2 DKO), a key downstream node in the Hippo pathway, displayed similar rates of ECAR as compared to parental cells following glucose exposure (Fig. 6C). However, the ability of FSK to suppress ECAR was significantly abrogated in the MST1/2 DKO cells, suggesting that MST1/2 and not the remainder of the Hippo pathway are important in this effect (Figs. 6C/D).

To determine whether MST1 and MST2 (encoded by the gene *Stk3* and *Stk4*) are involved in Epi-mediated p110α regulation *in vivo*, we generated mice with hepatocyte-specific genetic deletion of *Stk3* and *Stk4* (*Alb-Cre*^+/-^; *Stk3*^f/f^; *Stk4*^f/f^). These mice had significantly larger livers compared to their littermates without the *Alb-Cre* transgene, consistent with previous reports of hepatocyte hypertrophy following loss of MST1 and MST2 (Fig. S9D)(Zhou et al., 2009). Loss of MST1 and MST2 in hepatocytes resulted in a significant decrease in the phosphorylation of several target proteins including LATS1 (T1079) and MOB1 (T35) (Figs. 6E and S9E). We subjected these mice to acute Epi exposure via IP injection to assess the impact of these signaling changes on pT1061. While Epi increased the phosphorylation of T1061 in WT mice, this effect was significantly reduced in liver specific MST1/2 DKO mice (Figs. 6E/F). Interestingly, these mice also displayed increased AKT phosphorylation (pS473) following Epi, consistent with deletion of an endogenous negative regulator of p110α. We confirmed these results by inducing the deletion of MST1 and MST2 in adult mice by injecting an adenovirus containing Cre recombinase (AdCre) into *Stk3*^f/f^;*Stk4*^f/f^ mice (Fig. S9F/G).

Lastly, we evaluated the role of MST1/2 in the hepatic response to endogenous Epi by fasting the MST1/2 DKO mice and their littermate controls (*Stk3*^f/f^;*Stk4*^f/f^ referred to as Flox). Fasting increased the phosphorylation of p110α in Flox mice as compared to those that were “Fed,” however this change did not occur in DKO mice. We also interrogated the abundance of phosphorylated glycogen synthase (pGS) at site S641, an inhibitory modification controlled by PI3K signaling(Sakamoto et al., 2002). While the Flox mice could appropriately phosphorylate GS and suppress GS activity during fasting, these effects were absent in DKO mice (Figs. 7A-C). The activity of glycogen phosphorylase in the liver during fasting and rate of glycogenolysis following Epi exposure during a hyperinsulinemic euglycemic clamp were unchanged (Figs. 7D and S10). Together, the high glycogen synthase activity and normal glycogen phosphorylase activity associated with more glycogen content in the DKO livers during fasting (Fig. 7E). These data suggest that Epi acts via MST1/2 to suppress hepatic glycogen synthesis independent of glycogen phosphorylase activity(Petersen et al., 1998).

**Fig. 7.**
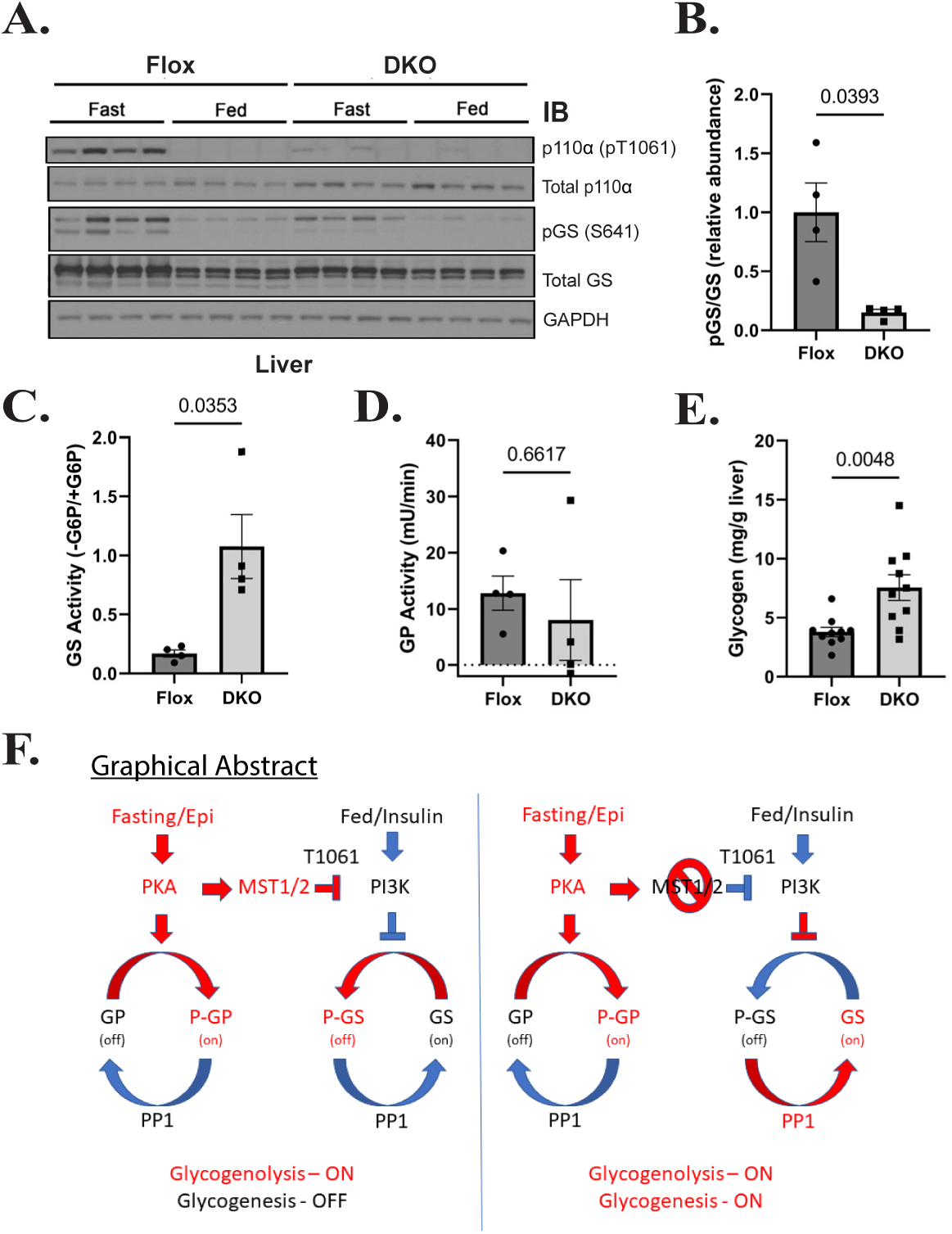
Loss of MST1/2 alters fasting glycogen metabolism. (A) Immunoblot for the indicated proteins using lysates from livers taken from Stk3^f/f^,Stk4^f/f^, Alb-Cre ^-/-^ (Flox) and Stk3^f/f^; Stk4^f/f^, Alb-Cre^+/-^ (DKO) that were fasted for 18 h (Fast) and then euthanized or provided food for 4 h (Fed). N=4. (B) Quantification of the ratio of phosphorylated glycogen synthase (pGS) to total glycogen synthase (GS) using band intensity for the Fasted mice in A. The ratio was normalized to the average of the Fasted Flox mice. Hepatic GS activity (C, N=4), glycogen phosphorylase (GP) activity (D, N=4), and glycogen content (E, N=10) were measured from Fasted Flox and DKO mice. Data is presented as Mean ± SEMs. Comparisons made via student’s t-test. (F) Schematic model depicting the normal fasting-induced regulation of glycogen metabolism including the activation of MST1/2 activity and subsequent phosphorylation and inhibition of PI3Kα. Loss of MST1/2 activity leads to persistent GS activity during fasting, likely via persistent PP1 activity.

## DISCUSSION

This is the first reported evidence that a signaling enzyme can inhibit p110α via direct modification and reveals an unexpected connection between the PI3K and Hippo pathways. Following phosphorylation by MST1/2, p110α undergoes a conformational change that prevents membrane binding and inhibits PIP_2_ phosphorylation. The p110α C-terminus is known to be essential for its function and harbors one of the most prevalent oncogenic point mutation of all cancers (H1047R), that conversely enhances membrane binding to promote PIP_2_ phosphorylation. Phosphorylation of p110α by MST1/2 provides cells the opportunity to regulate this process, and like the H1047R mutation, this effect is alpha isoform specific. Our work also implicates p110α as an essential mediator and regulatory control point linking cell growth pathways with those regulating cell metabolism.

Several independent laboratories have observed the development of insulin resistance in humans exposed to Epi(Bessey et al., 1983; Deibert and DeFronzo, 1980), but it has remained unclear how PI3K signaling is dampened in this setting. Our data show that Epi results in PKA-mediated activation of MST1/2, which then phosphorylate and inhibit p110α. These data are in-line with those of Liu *et al*. who demonstrated that hepatic MST1/2 is sequestered in glycogen via an interaction with Laforin(Liu et al., 2021). Together, our results support a model where counter-regulatory hormones regulate hepatic insulin signaling by activation of PKA-mediated glycogen degradation, MST1/2 release, and direct phosphorylation and inhibition of p110α. In this setting, MST1/2 is also freed to phosphorylate LATS1/2 and reduce the abundance of YAP/TAZ in hepatocytes. Low levels of TAZ contribute to the activation of gluconeogenesis in the liver via interactions with the glucocorticoid receptor(Xu et al., 2021).

Our study focused on the acute changes in hepatic PI3K signaling following administration of Epi. Glycogen synthesis and the downstream phosphorylation events in the PI3K pathway are the most direct readouts of hepatocellular insulin action(Petersen and Shulman, 2018). In the liver, insulin promotes glycogen synthesis and inhibits glycogenolysis to suppress hepatic glucose production(Lewis et al., 2021). In contrast, Epi increases hepatic glucose release by activating glycogenolysis and inhibiting glycogen synthase(Cori and Cori, 1929; Jensen et al., 2011). The loss of MST1/2 in the liver creates a unique scenario where Epi continues to activate glycogenolysis, but the crosstalk to PI3K is lost and glycogen synthesis continues.

The long-term implications of this pathway on glucose homeostasis remain unclear. It is interesting to note that other GCK family kinases have been reported as negative regulators of insulin sensitivity. For example, the overexpression of YSK1/STK25 reduces glucose tolerance and insulin sensitivity in mice fed a HFD(Cansby et al., 2013), the deletion of MST3/STK24 improves hyperglycemia and insulin resistance(Iglesias et al., 2017), and the deletion of HGK improves insulin response in adipocytes(Virbasius and Czech, 2016). Therefore, the GCK kinases could be an attractive therapeutic target to improve glycemia.

## ACKNOWLEDGMENT

We thank Ismail Moarefi and Karine Roewer of CreLux WuXi AppTec for assisting in solving the structure of the p110α(T1061E)/ni-p85α complex. We thank David McElliot of Petra Pharmaceuticals for helping coordinate our work with CreLux. We thank Kun-Liang Guan for providing CRISPR-engineered HEK293 cells. Isolation of primary hepatocytes was performed by the Metabolic Phenotyping Core at Weill Cornell Medicine. Euglycemic clamp studies were performed by the Rodent Metabolic Phenotyping Core supported by Penn Diabetes Research Center grant P30-DK1952, and the Vanderbilt Mouse Metabolic Phenotyping Center (DK059637). The Vanderbilt Hormone Assay and Analytical Core performed the insulin and catecholamine analyses (DK059637 and DK020593). This work was supported in part by NIH CA230318 (M.D.G.), NIH R35 CA197588 (L.C.C.), a grant from the Breast Cancer Research Foundation (L.C.C.)

## AUTHOR CONTRIBUTIONS

T.Y.L, J.L.J, L.C.C and M.D.G. conceived and designed the study. T.Y.L, K.L, J.L.J performed the protein purification, molecular cloning and liposome sedimentations assays. T.Y.L and J.L.J performed the thermal shift assays. J.L.J performed the activity screens, lipid kinase activity assay, peptide library assays, and structural analyses. N.V., K.K., and E.A.K. provided structure insights. N.V. and K.K. performed the ADP-Glo assays. T.M.Y. performed computational analyses on the peptide library data. T.W. produced the phosphor-specific polyclonal antibody for this work. G.Z. and M.Z. performed mass spectrometry analyses. Y.M. provided technical advances to the protein purification. T.Y.L, S.R, S.K.H, K.L, E.R.K and B.D.H performed the mouse experiments. T.Y.L, S.K.H and K.L performed the immunoblotting. T.Y.L performed the cultural assays. R.E.S isolated and cultured human hepatocytes. M.D.G. performed the seahorse assay. T.Y.L, M.D.G, J.L.J and M.N.P. performed the data analysis. T.Y.L, J.L.J, and M.D.G. wrote the manuscript. All authors assisted with data interpretation and contributed to the editing of the manuscript.

## MATERIALS and METHODS

### Recombinant kinases

Recombinant kinases MST1, HGK, MST2, MST3, TAOK1, GLK, AKT1, AMPK1, ERK2, PDK1, CK1A, GSK3A, IKKB, IRAK4, ASK1, GCN2, EEF2K, JAK1, CHK1, CHK2, DAPK1, PKACA, PKCA, PAK1, AurB, CDK1, DYRK1A, SRC, and ABL1 were purchased from SignalChem. Recombinant kinases ATM, LKB1, and TAK1 were purchased from Millipore. Recombinant kinases LATS2 and OXSR1 were purchased from Abnova. Recombinant CK2 was purchased from NEB. Recombinant TGFBR1 was purchased from Proqinase. Recombinant kinase WEE1 was purchased from BPS Biosciences. Recombinant PIK3CG was purchased from Thermo Fisher.

### Mutagenesis and cloning

pBabe puro HA-PIK3CA was purchased from Addgene (plasmid #12522). This construct has an artifactual amino acid change in its coding sequence (I143V) and site-directed mutagenesis (Quikchange II XL, Agilent) was performed to convert it back to WT with following two primers (all primers were ordered from Integrated DNA Technologies):

*PIK3CA* V143I to WT isoleucine Forward: GACTTCCGAAGAAATATTCTGAACGTTTGTAAA
*PIK3CA* V143I to WT isoleucine Reverse: TTTACAAACGTTCAGAATATTTCTTCGGAAGTC.

The coding sequence of PIK3CA was then cloned *PIK3CA* into the pcDNA3.4 vector through in-fusion cloning (Clontech) as an untagged protein or with an N-terminal polyhistidine tag. Sequencing analysis was performed with Snapgene.

### *PIK3CA* T1061A mutagenesis primers

*PIK3CA* T1061A_FW: 5’-caaaaatggattggatcttccacgcaattaaacagcatgcattgaac-3’
*PIK3CA* T1061A_RV: 5’-gttcaatgcatgctgtttaattgcgtggaagatccaatccatttttg-3’

### *PIK3CA* T1061E mutagenesis primers

*PIK3CA* T1061E_FW: 5’-cctcagttcaatgcatgctgtttaatctcgtggaagatccaatccatttttgttg-3’
*PIK3CA* T1061E_RV: 5’-caacaaaaatggattggatcttccacgagattaaacagcatgcattgaactgagg-3’

### *PIK3CA* W1057A mutagenesis primers

*PIK3CA* W1057A_FW: 5’-ctgtttaattgtgtggaagatcgcatccatttttgttgtccagcca-3’
*PIK3CA* W1057A_RV: 5’-tggctggacaacaaaaatggatgcgatcttccacacaattaaacag-3’

### Protein purification

Expi293F cells (Thermo Fisher) were cultured in Expi293 Expression Medium (Thermo Fisher) and incubated in 2L spinner flasks on an orbital shaker at 90 rpm at 37 °C with 8% CO_2_. 350 μg of pcDNA 3.4-PIK3CA and 150 μg pcDNA 3.4-His_6_-flag-TEV-PIK3R1 were combined and diluted in Opti-MEM I Reduced Serum Medium (Thermo Fisher). ExpiFectamine 293 Reagent (Thermo Fisher) was diluted with Opti-MEM separately then combined with diluted plasmid DNA for 10 minutes at room temperature. The mixture was then transferred to 500 mL EXPI-293F cells (3 × 10^6^ cells/mL) and incubated. After 24 hours, ExpiFectamine 293 Transfection Enhancer 1 and Enhancer 2 (Thermo Fisher) were added. Two days later, cells were centrifuged at 300 × g for 5 min, snap freezed in liquid nitrogen and store at -80 °C (3 days post-transfection). All steps of protein purification were performed in the cold room at 4°C. Cell pellets were solubilized in lysis buffer (50 mM Tris pH 8.0, 400 mM NaCl, 2 mM MgCl_2_, 5% glycerol, 1% Triton X-100, 5 mM β-mercaptoethanol) supplemented with EDTA-free phosphatase and protease inhibitor cocktail (Life technologies) and lysed by Dounce homogenization (20 strokes). Lysates were centrifuged at 100,000 x g for 1 h and the supernatant were affinity purified on Ni-NTA resin (Qiagen) by batch binding for 30 min-1h. Resin was washed with 10 column volumes of base buffer (50 mM Tris pH 8.0, 500 mM NaCl, 2 mM MgCl_2_, 2% glycerol, 20 mM imidazole) and eluted in 10 column volumes of elution buffer (50 mM Tris pH 8.0, 100 mM NaCl, 2 mM MgCl_2_, 2% glycerol, 250 mM imidazole). Eluted protein was concentrated using 30 kDa Ultra Centrifugal Filter Units (Amicon) and supplemented with 1 mM DTT and 25% glycerol before snap freezing in liquid nitrogen. The same approach was carried out for purification of untagged p110β: His_6_-p85 α and untagged p110δ: His_6_-p85α.

### Lipid kinase assay

For phosphorylation reactions, predetermined concentrations of protein kinase and PI3K complexes were added to master mix containing 50 μM of ATP, 0.1 mg/mL of BSA and 1x Assay Buffer I (SignalChem) in 40 μl total volumes. Reactions were carried out at 30 °C for 30 minutes. Next, 0.01 mCi/mL ^32^P-labeled ATP (Perkin Elmer) and 25 μM PIP_2_ (Thermo Scientific) were included, bringing total volume to 100 μL. PIP_2_ phosphorylation reactions were carried out at 30°C for 20 min. 50 ul of 4N HCl was added to quench the reaction and the lipid was extracted with 100 μl of CHCl3: MeOH (1:1) followed by vortexing for 1 minute and centrifugation at maximum speed for 2 minutes. 10 μL of the lower hydrophobic phase was extracted with gel loading pipet tips and spotted onto a silica plate (EMD Millipore #M116487001) for thin layer chromatography. Plates were placed in a sealed chamber with 1-propanol: 2M acetic acid (65:35). Radioactivity was imaged with a Typhoon FLA 7000 phosphorimager (GE) and quantified by ImageQuant (GE). For cationic liposomes, EPC (#890704C) and PI (840042C) were purchased from Avanti. Liposomes containing 5% PI, 60% EPC, and 35% cholesterol were prepared by extrusion via 20 passes through a 0.1 μm membrane using a Mini-Extruder kit (Avanti). The anionic liposomes in these experiments replaced EPC with PS.

### Thermal shift assays

500 nM p110α/p85α -/+ 100 nM MST1, was added to master mix containing 50 μM of ATP, 0.1 mg/mL of BSA and 1x Assay Buffer I (SignalChem) to a total volume of 50 μL into a MicroAmp Optical 8-Cap strip (Thermo Fisher). Tubes were placed in a Thermocycler (BioRad). Samples were cycled at 50 °C for 30 seconds, then on a temperature gradient from 50 °C – 65 °C for 3 minutes, then 25 °C for 3 minutes. Condensates were collected by minispin for 30 seconds and 40 μL of the supernatant was transferred to separate Eppendorf tubes. Tubes were spun at maximum speed in a microfuge for 20 minutes at 4 °C. 30 μL of the supernatant was transferred to separate Eppendorf tubes and 10 μL of 4× LDS Sample Buffer were added. Samples were run out on 4%-12% Bis-Tris Pre-cast Gel (Life Technologies) and soluble p110α was probed by Western blotting across the temperature gradient using anti-p110α antibody.

### Liposome preparation and liposome sedimentation assays

PS (#840032) and PE (#840026C) were purchased from Avanti and cholesterol (#CH800) was purchased from Nu Chek Prep. Lipid stocks were prepared at 35% PE, 30% PS, and 35% cholesterol. N_2_ gas was applied for 15 seconds and tubes were frozen and stored at -20 °C. Before the experiments, the lipid stocks were vortexed and 100 μL of chloroform (HPLC-grade) was transferred to a clean glass vial. N_2_ gas was immediately applied to the stock tube, capped, and stored at -20 °C. N_2_ gas was applied to the 100 μL aliquot to evaporate chloroform. Next, 1 mL of 20 mM HEPES/ 1 mM EGTA was added and lipids were rehydrated at room temperature for 1 h. Liposomes were prepared by extrusion via 7 passes through a 0.8 μm membrane using a Mini-Extruder kit (Avanti). Liposomes were transferred to a clean Eppendorf tube and centrifuged at 10,000 × g for 5 minutes. Supernatant was discarded, and the lipid pellet was resuspended in 50 μL 1× Assay Buffer I (SignalChem), vigorously until resuspended. In parallel, p110α phosphorylation assays were carried out. 150 μL volume of reaction mix, containing 500 nM p110α/p85α, -/+ 100 nM MST1, 50 μM of ATP, 0.1 mg/mL of BSA and 1× Assay Buffer I (SignalChem) were added to liposomes and 50 μL volumes were distributed across three Eppendorf tubes. Binding reactions proceeded for 10 minutes at RT. Samples were centrifuged at 10,000 × g for 5 min at RT and 40 μL supernatant was carefully removed without disturbing the pellet and optimal amount of NuPAGE 4× LDS Sample Buffer (Life Technologies #NP0008) was added. Lipid pellets were resuspended with 40 μL of Buffer I with optimal amount of 4× LDS Sample Buffer added. The amount of membrane-bound p110α was probed and analyzed by Western blotting. For quantification, densitometry was performed using ImageStudioLite.

### X-ray crystallography

The constructs used for crystallization comprise residues 1 to 1068 of the catalytic subunit p110 with the phosphomimetic mutation T1061E and residues 308-593 of the regulatory subunit p85. p110α was expressed in insect cells (SF21) as a complex with p85α. Crystals of the p110α (T1061E) /p85 heterodimer in complex with GDC-0077 were obtained using hanging-drop vapor-diffusion set-ups. The p110α (T1061E)/ p85 complex at a concentration of 9 mg/ml (20 mM Tris/HCl, 150 mM NaCl, 1 mM TCEP, pH 8.0) was pre-incubated with 0.5 mM (6.9-fold molar excess) of GDC-0077 (10 mM in DMSO) for 1 h. 1μL of the protein solution was then mixed with 1 μL of reservoir solution (0.10 M Bis-Tris-Propane pH 8.10, 0.20 M Na 3 -citrate, 10.00 %(w/v) PEG 3350) and equilibrated at 20 °C over 0.4 ml of reservoir solution. Well diffracting crystals appeared within 4 days and grew to full size over 20 days. PI3K structural display and mapping was performed using PyMOL.

### Transcreener assay

Transcreener ADP fluorescence intensity assay (Bellbrook Labs) was applied to determine the ATPase activity of the PI3K complex under different treatment. For preparing the HGK-treated PI3K phosphorylation reaction, 38 μL of PI3K complex (1.3 mg/ml) (EMD Millipore #14-602M) was incubated with 30 μL of HGK, 20 μL of ATP (1 mM) and 412 μL of kinase Buffer to the total volume of 500 μL. Reaction was carried out at 30 °C for 1hr. GST-HGK was removed from the reaction using 10 μL of Glutathione Sepharose beads (Thermo Fisher Scientific #45000285) and batched incubation at 4 °C for 30 min. The PIK3CA proteins were diluted to 5X stocks in buffer (20 mM HEPES pH 7.4, 100 mM NaCl, 0.5mM EGTA, 0.01% triton-x-100) just before use. 10mM stocks of Alpelisib were serially diluted (3×) in neat DMSO and stored in a dessicator at room temperature. PIK3CA, along with buffer components (except ATP) were incubated with or without compound at 27 °C for 1h. After incubation, the reaction was initiated by the addition of 5uL of 125uM ATP. A typical assay mixture (25 uL) contained 40mM HEPES buffer, pH 7.4, 25 mM MgCl2, 0.01% v/v triton-X-100, 5% v/v DMSO, 20 mM NaCl, 1-5 nM Wt or H1047R, 25 uM ATP, and 50 uM PIP2diC8 or PIP2 in membrane. The reaction was allowed to proceed till ∼10% conversion (2.5 uM ADP) after which time, 10 uL of reaction mixture was quenched with 25uL of transcreener reagent (transcreener ADP2 FI assay kit, BellBrook labs, Cat. No. 3013). The contents were incubated at RT for 1h and fluorescence was measured using a plate reader (Paradigm, Molecular Devices). The same assay was also run at pH 6.0 or 6.4 using MOPS buffer. A calibration curve was generated under identical buffer conditions with varying ADP amounts. Using that, the observed fluorescence was converted to uM ADP. A plot between [ADP] and log[I] yielded the dose-response curves that enabled the calculation of IC_50_.

### LC-MS/MS-based phospho-peptide detection and analyses

MST1-treated and untreated PI3K phosphorylation reaction were prepared as followed: 2 μL of PI3K complex was incubated -/+ 4.5 μL MST1 and the mastermix containing 2 μL of ATP (1 mM) and 31.5 μL of kinase Buffer to the total volume of 40 μL. Reaction was carried out at 30 °C for 1h and stopped by adding 4x sample buffer. Samples were running on the SDS-PAGE and gel was stained using QC Colloidal Coomassie Stain (Bio-Rad Laboratories #1610803). The sample was digested in-gel with trypsin overnight at 37 °C following reduction with DTT and alkylation with iodoacetamide. The digest was vacuum centrifuged to near dryness and desalted by C18 stage-tip column. A Thermo Fisher Scientific EASY-nLC 1000 coupled on-line to a Fusion Lumos mass spectrometer (Thermo Fisher Scientific) was used. The raw files were processed using the MaxQuant(Cox and Mann, 2008) computational proteomics platform version 1.5.5.1 (Max Planck Institute, Munich, Germany) for protein identification. The fragmentation spectra were used to search the UniProt human protein database (downloaded September 21, 2017). Oxidation of methionine, protein N-terminal acetylation and phosphorylation of serine, threonine and tyrosine were used as variable modifications for database searching. The precursor and fragment mass tolerances were set to 7 and 20 ppm, respectively. Both peptide and protein identifications were filtered at 1% false discovery rate based on decoy search using a database with the protein sequences reversed. Identified phosphopeptides were subjected to manual inspection for confirmation of the identification.

### Peptide library arrays and computational prediction of phosphorylation

Recombinant MST1 was distributed across 384-well plate, mixed with the peptide substrate library (Anaspec #AS-62017-1 and #AS-62335), kinase assay buffer I (SignalChem) and γ-^32^P-ATP, and incubated for 90 mins at 30°C. Each well contains a mixture of peptides with a centralized phosphor-acceptor (serine and threonine at a 1:1 ratio), one fixed amino acid in a randomized background. All 20 amino acids were distributed across the range of -5 to +4, relative to the centralized serine/threonine to determine the individual contributions of amino acids along the substrate peptide, as shown in the heatmap where the x-axis represents fixed amino acid and the y-axis represents the relative position. Peptides contained C-terminal biotinylations and were spotted on streptavidin-conjugated membranes (Promega #V2861) and imaged with Typhoon FLA 7000 phosphorimager (GE) and quantified by ImageQuant (GE). Intensities were normalized by position to generate a PSSM for MST1. MST1’s PSSM was normalized position-wise (dividing individual amino acid intensities at position -5 by the average intensity at this position, then position -4, etc.) and utilized to score threonines in p110α’s amino acid sequence. The products of their corresponding values at positions -5 to +4 determined their final scores.

### pT1061 peptide phosphorylation assays

A synthetic peptide modeled after p110α’s C-terminus (sequence: [K(LC-biotin)]-K-G-A-<u>M-D-W-I-F-T-I-K-Q-H-A-L-N, where LC=linker chain</u>) were incubated with reaction mixtures, kinase assay buffer I (SignalChem) and γ-^32^P-ATP, for 5 min at 30°C. Reactions were spotted onto streptavidin-conjugated membranes (Promega #V2861) and imaged with Typhoon FLA 7000 phosphorimager (GE).

### Illustrations

Experimental schema and illustrative models were generated by BioRender (https://biorender.com/) and Chemdraw. Sequence alignments performed with Geneious. Kinome images generated and modified at http://kinhub.org/kinmap/index.html.

### Cell culture and cell Lines

All cell lines were cultured at 37 °C with 5% CO_2_. MDA-MB-231, Huh7, HEK293A cell lines were cultured in DMEM (Life Technologies #11965092) supplemented with 10% FBS. MCF10A were cultured in DMEM/F12 (Life Technologies #11330032) containing 5% horse serum, 20 ng/ml EGF, 0.5 µg/ml hydrocortisone, 10 µg/ml insulin, 100 ng/ml cholera toxin, and 50 µg/ml penicillin/streptomycin (P/S). Primary mouse hepatocytes were cultured in William’s E medium (Life Technologies #12551032) containing 10% FBS and 1% P/S. AML12 cells were cultured in DMEM: F12 medium (ATCC #30-2006) supplemented with 10% FBS, Insulin-Transferrin-Selenium (ITS-G) (100X) (Thermo Fisher Scientific #41400045), and 40 ng/ml dexamethasone. For signaling assays, cells were washed 1x in PBS and placed in starvation media (-FBS) for 2-16 h depending upon cell line.

### Immunoprecipitation and lipid kinase assay

Cell were lysed using the PI3K buffer [25 mM Tris, pH 7.5, 10 mM EDTA, 10 mM EGTA, 1% Nonidet P-40] with one tablet (per 10 mL) of protease and phosphatase inhibitor (Life Technologies). After lysis on ice for 30 min, cell lysates were centrifuged for 10 min, 4 °C and supernatant were used for immunoprecipitation. Lysates were incubated with anti-phospho-Tyr-4G10 (EMD Millipore 05-321), anti-p110α (CST #4249) or anti-p85α antibody (EMD Millipore #ABS233) for 1 h, followed by protein A/G beads (Santa Cruz) for additional 1hr, and then washed twice with PI3K buffer, and three times with TNE buffer [10 mM Tris (pH 7.5), 100 mM NaCl, 1 mM EDTA]. For PI3K activity assays, the beads (50 μL) were incubated in 1× Assay Buffer I (SignalChem), 0.5 μL ATP (10 mM), 1 μL ^32^P-labeled ATP (0.01 mCi/mL) and 10 μL of PIP_2_ (250 μM) in 100 μL total volumes. Reactions were carried out at 30°C for 30 min. 50 ul of 4N HCl was added to quench the reaction and the lipid was extracted with 100 μl of CHCl3: MeOH (1:1) followed by vortexing and centrifugation. 20 μL of the lower hydrophobic phase was extracted with gel loading pipet tips and spotted onto a silica plate (EMD Millipore, M116487001) for thin layer chromatography. Plates were placed in a sealed chamber with 1-propanol: 2M acetic acid (65:35). Radioactivity was visualized and quantified using a Typhoon FLA 7000 phosphorimager.

### Immunoblotting

Cell lysates were prepared in the cell lysis buffer [50 mM Tris-HCl [pH 7.4], 150 mM NaCl, 1 mM EDTA and 1% NP-40 with one tablet (per 10 mL) of protease and phosphatase inhibitor (Life Technologies)]. After incubation on ice for 30 min, lysates were centrifuged at top speed for 10 min and supernatant were collected and proteins were quantified using Protein DC assay (BioRad). Proteins were running and separated on 4%-12% Bis-Tris Pre-cast Gel (Life Technologies) using MOPS buffer and transferred to PVDF membrane at 350 mA for 1.5 h. Membranes were blocked in 5% non-fat milk in TBST and incubated with primary antibody overnight. Primary antibody against pAKT (S473) [#3787, #4060], pAKT (T308) [#4056], pAKT2 (S474) [#8599], pan AKT [#2920], AKT1 [#2967], AKT2 [#3063], pCREB (S133) [#9198], pLATS1 (T1079) [#8654], pMOB1 (T35) [#8699], pYAP (S127) [#13008], pPKA substrate motif antibody [#9624], p110α [#4249], CREB [#4820], MOB1 [#13730], LATS1 [#9153], MST1 [#14946] and GAPDH [#5174] were from Cell Signaling Technology. MST2 [#ab52641] was from Abcam, and p110 (pT1061) polyclonal antibody was developed and purified by Cell Signaling Technology. HRP-conjugated secondary antibody was used at 1:5000 in 5% milk and membrane were developed using ECL solution and exposed to film. 7.5% phos-tag gel was used to resolve phosphor-p110α (T1061) (Wako Diagnostics/Chemicals # 192-18001).

### Vectors and compounds

Alpelisib (MEDCHEM EXPRESS LLC #HY-15244), Neratinib (Selleck #S2150), Go6976 (EMD Millipore #365250), DMX-5804 (MEDCHEM EXPRESS LLC# HY-111754), XMU-MP-1 (Selleck #S8334), H-89 (Tocris Bioscience #2910), AT13148 (Selleck #S7563) and Propanolol HCl (Sigma Aldrich #P0884) were purchased and reconstituted using DMSO and stored at -20 before use. cDNA of PKIA were synthesized by IDT (gBlock) and PCR amplified using the two primers: pLenti-PKIA-IF-FW: 5’CGACTCTAGAGGATCCATGACTGATGTGGAAACTACATATG; pLenti-PKIA-IF-Rev: 5’-GAGGTTGATTGTCGACTTAGCTTTCAGATTTTGCTGCTTC and PCR products were purified. Vector pLenti-GFP were double digested with BamHI-HF and SalI-HF and gel purified. Next, PCR product were inserted into the vector backbone using In-fusion HD cloning (Takara) and transformed in Stbl3 competent cells and grown at 37°C overnight.

### Mouse strain and breeding

All animal studies were conducted following IACUC approved animal protocols (#2013-0116) at Weill Cornell Medicine. Mice were maintained in temperature- and humidity-controlled on a 12-h light/dark cycle and received a normal chow diet with free access to drinking water. C57/BL6 mice were purchased at 8 weeks of age from Jackson laboratories (Bar Harbour, ME). *Stk3*^f/f^, *Stk4*^f/f^ mice were purchased from Jackson laboratories (Stock No: 017635). Liver specific MST1/2 double knockout mice (*Stk3*^f/f^, *Stk4*^f/f^, *Alb-Cre*^-/+^) were generated by breeding *Stk3*^f/f^, *Stk4*^f/f^ mice with Albumin-Cre (Stock No: 003574) mice. Homozygous null mice for the *Adrb1* and *Adrb2* genes (*Adrb1*^-/-^; *Adrb2*^-/-^)were purchased from the Jackson laboratory (Stock No. 003810). Adipocyte-specific deletion of *Pnpla2* gene were generated by breeding *Pnpla2*^f/f^ mice with *Adipoq*-*Cre* mice. *Pnpla2*^f/f^ mice were purchased from the Jackson laboratory (Stock No.024278). All mice were fed normal chow (Purina 5053) or a 60% high-fat diet (D12492i, Research Diets, Inc. New Brunswick, NJ, USA), when indicated.

### Mouse injections and metabolic tissue collections

Fed male mice at 10 weeks of age received an intraperitoneal injection of epinephrine (0.3 μg/g) or insulin (0.75 mU/g body weight) for the indicated durations. For assessment of blood glucose, 10 μl of blood was taken from the tail of mice before treatment and at the indicated time points using a handheld glucose meter (OneTouch). Mice were sacrificed using cervical dislocation and the metabolic tissues were harvested and immediately frozen in liquid nitrogen. Samples were stored in -80 °C until use. Frozen tissues were homogenized in 1 ml of lysis buffer (50 mM Tris·HCl (pH 7.4), 150 mM NaCl, 1 mM EDTA, 10% glycerol, 1% Nonidet P-40, 0.5% Triton X-100) and 1 tablet (per 10 mL) of protease and phosphatase inhibitor (Life Technologies). Homogenates were centrifuged at top speed for 30 min and 50 μg of protein, determined by Protein DC Assay (BioRad) were used for Western blot analysis.

### Primary hepatocytes isolation

Mouse hepatocytes were isolated by a 2-step perfusion procedure(Scapa et al., 2008). Briefly, mice were anesthetized with ketamine (100 mg/kg), xylazine (10 mg/kg), and acepromazine (3 mg/kg). Following laparotomy, the liver was perfused *in situ* through the inferior vena cava with 20 ml of prewarmed liver perfusion medium (Invitrogen) followed by 40 ml of liver digestion medium (Invitrogen). The liver was then placed in ice-cold hepatocyte wash medium (Invitrogen), and the capsule of the liver was then gently disrupted in order to release the hepatocytes. The cell suspension was filtered with a 70-μm cell strainer (Becton, Dickinson), and cells were washed once (30× g, 4 min, 4°C). Dead cells were removed using Percoll solution (Sigma-Aldrich) as described previously(Lee et al., 2004). Live cells were washed, pelleted, resuspended in incubation medium, and seeded on 6-well Primaria plates (Becton, Dickinson) at a density of 4 x10^5^ cells/ well. Cells were allowed to adhere to the plates for 3 h, after which the culture medium were replaced by fresh 2 ml medium.

### Extracellular acidification rates measurement (ECAR)

Cells were plated in XF96 V3 PET cell culture microplates (Seahorse Biosciences, Cat. #; 101104-004) at a density of 20,000 cells per well in DMEM (Life Technologies #11965092) with 10% FBS. After 24 h, growth media was changed to bicarbonate-free assay media (XF assay medium, Seahorse Biosciences, Cat. #: 102365-100) and incubated at 37**°**C for 1 h in a CO_2_-free incubator. Extracellular acidification rate (ECAR) was measured using an XF96 Extracellular Flux Analyzer (Seahorse Biosciences).

### Hyperinsulinemic-euglycemic clamps

#### Rat study

All procedures for the rat hyperinsulinemic–euglycemic clamp were approved by the University of Pennsylvania Animal Care and Use Committee. Rats were fasted for 5 hours prior to the initiation of the clamp and acclimated to the bucket containers. Jugular vein and carotid arterial line were hooked up to the dual swivel 2 h prior to the clamp initiation. During the 90 min clamp period, rats received a constant infusion of insulin (Novolin Regular) at 5 mU/kg/min. Blood glucose levels were clamped at euglycemia with a variable infusion of 50% dextrose. Blood glucose levels were monitored via arterial blood sampling every 10 minutes and glucose infusion rates adjusted to maintain euglycemia. 87 minutes after initiation of the clamp, rats received either an IV infusion of vehicle saline for 10 min or epinephrine 0.75 ug/kg/min for 10 or 30 min. Blood samples were collected at the beginning of the clamp (−30 and 0 min, basal period) and during the clamp period (30, 60, 90, 100 or 120 min) for the subsequent assay of catecholamines and insulin levels. At study termination, animals were anesthetized with IV pentobarbital and tissues collected and flash frozen in liquid nitrogen for livers.

#### Mouse study

All procedures for the hyperinsulinemic–euglycemic clamp were approved by the Vanderbilt University Animal Care and Use Committee. Catheters were implanted into a carotid artery and a jugular vein of mice for sampling and infusions respectively five days before the study as described by Berglund et al(Berglund et al., 2008). Insulin clamps were performed on 5 hr-fasted conscious unhandled mice, using a modification of the method described by Ayala et al(Ayala et al., 2006). After 3h of fast, an arterial blood sample was obtained to determine natural isotopic enrichment of plasma glucose. Immediately following this sample, a quantitative stable isotope delivery to increase isotopic enrichment above natural isotopic labelling was initiated as described previously(Hasenour et al., 2015). Briefly, a [6,6-^2^H_2_]Glucose-^2^H_2_O (99.9%)-saline bolus was infused for 25 min to enrich total body water to 4.5% (t=-120 min to -95 min). A continuous infusion of [6,6-^2^H_2_]glucose (t=-95min to 0min; 0.8 mg kg^−1^ min^−1^) was started following the [6,6-^2^H_2_]Glucose-^2^H_2_O-saline prime. The insulin clamp was initiated at t=0min with a continuous insulin infusion (2.5 mU/kg body weight/min). At the same time, a variable infusion of glucose was started (50% dextrose + ^2^H_2_O (0.04 MPE) +[6,6-^2^H_2_]Glucose (0.08 MPE)) in order to maintain stable euglycemia and stable enrichment of ^2^H_2_O and [6,6-^2^H_2_]Glucose in plasma. At t=91min, an epinephrine infusion was initiated (continuous infusion until sacrifice; 4 ug/kg body weight/min, in 0.7 mg/mL ascorbate for stability). Washed red blood cells were also continuously infused during the clamp period to maintain hematocrit. Every infusate was prepared in a 4.5% ^2^H _2_O - enriched saline solution. Arterial glucose levels were monitored every 10 min to provide feedback for adjustment of the glucose infusion rate (GIR). Blood sampling for glucose kinetics was performed at t=- 10;-5 (basal), t=80-90 (insulin) and t=110-140 min (insulin+epinephrine) of the clamp. Clamp insulin was determined at *t*=120 min. At 140 min, 13µCi of 2[^14^C]deoxyglucose ([^14^C]2DG) was administered as an intravenous bolus. Blood was taken from t=142-165min for determination of plasma [^14^C]2DG. Plasma lactate and catecholamines were determined at t=165min. At t=166min, mice were sacrificed by pentobarbital injection, and tissues immediately frozen in liquid nitrogen. Plasma glucose enrichments ([6,6-^2^H_2_]Glucose), isotopomer distribution and the enrichment ratio of deuterium on the fifth (C5) and second carbon (C2) of glucose were assessed by GC-MS as described previously(Hughey et al., 2017). Glucose fluxes were assessed using non–steady-state equations (volume of distribution of glucose= 130 ml/kg)(Steele et al., 1956). The contribution of gluconeogenesis was assessed as the ratio of C5 and C2 of plasma glucose(Antoniewicz et al., 2011; Burgess et al., 2004). [^14^C]2DG in plasma samples, and [^14^C]2DG-6-phosphate in tissue samples were determined by liquid scintillation counting. The glucose metabolic index (Rg) was calculated as previously described(Kraegen et al., 1985). Plasma insulin was determined by RIA. Plasma lactate was determined by colorimetric assay (abcam). Plasma catecholamine levels were determined by HPLC. Stable isotopes were purchased from Cambridge Isotope Laboratories, Inc. (Tewksbury, MA). Radioactive tracers were purchased from Perkin Elmer.

### Catecholamine measurements

Mice were sacrificed using cervical dislocation and blood were collected immediately using cardiac puncture. EGTA-glutathione solution (pH6.0-7.4) [4.5g EGTA and 3.0g Glutathione/ per 50 mL] was added at the time of blood collection (2 μl of solution was added per 100 μl of blood). After centrifuged at 4 °C, 10000 rpm for 10 min, supernatants were collected and aliquoted and samples for catecholamine analysis were flash frozen in liquid nitrogen. Plasma for insulin measurements were stored at -20 °C. Catecholamine and insulin measurements were performed by the VUMC Hormone Assay and Analytical Services Core.

### Glycogen synthase activity, phosphorylase activity, and tissue content

Glycogen synthase activity was measured using the incorporation of glucose from UDP-[6-^3^H] D-glucose into glycogen using liver homogenates as previously described(Li et al., 2019). Activity was calculated as the ratio of activated GS activity (-G6P) versus the total GS activity (+G6P). Glycogen phosphorylase activity was measured from liver homogenates using a calorimetric assay that measures the appearance of G1P in the presence of excess substrate (Abcam ab273271). For glycogen tissue content, frozen liver tissue (30–50 mg) and dilutions of glycogen type III obtained from rabbit liver (Sigma-Aldrich) were homogenized in 0.03 N HCl. An aliquot of the homogenate was mixed with 1.25 N HCl and heated for 1 h at 95 °C. Samples were centrifuged at 18,400 × g, and 10 µL of supernatant was mixed with 1 mL of glucose oxidase reagent (Stanbio Laboratory). After a short incubation at 37 °C, the absorbance was read at 505 nm.

### Statistics

All summary data are expressed as mean ± SEM. One-way, two-way ANOVA or student’s *t* test were used as indicated followed by correction for multiple comparisons test using Prism 7 (GraphPad La Jolla, CA). Statistical significance is indicated as specific p values in figures.

---

**Fig. S1.**
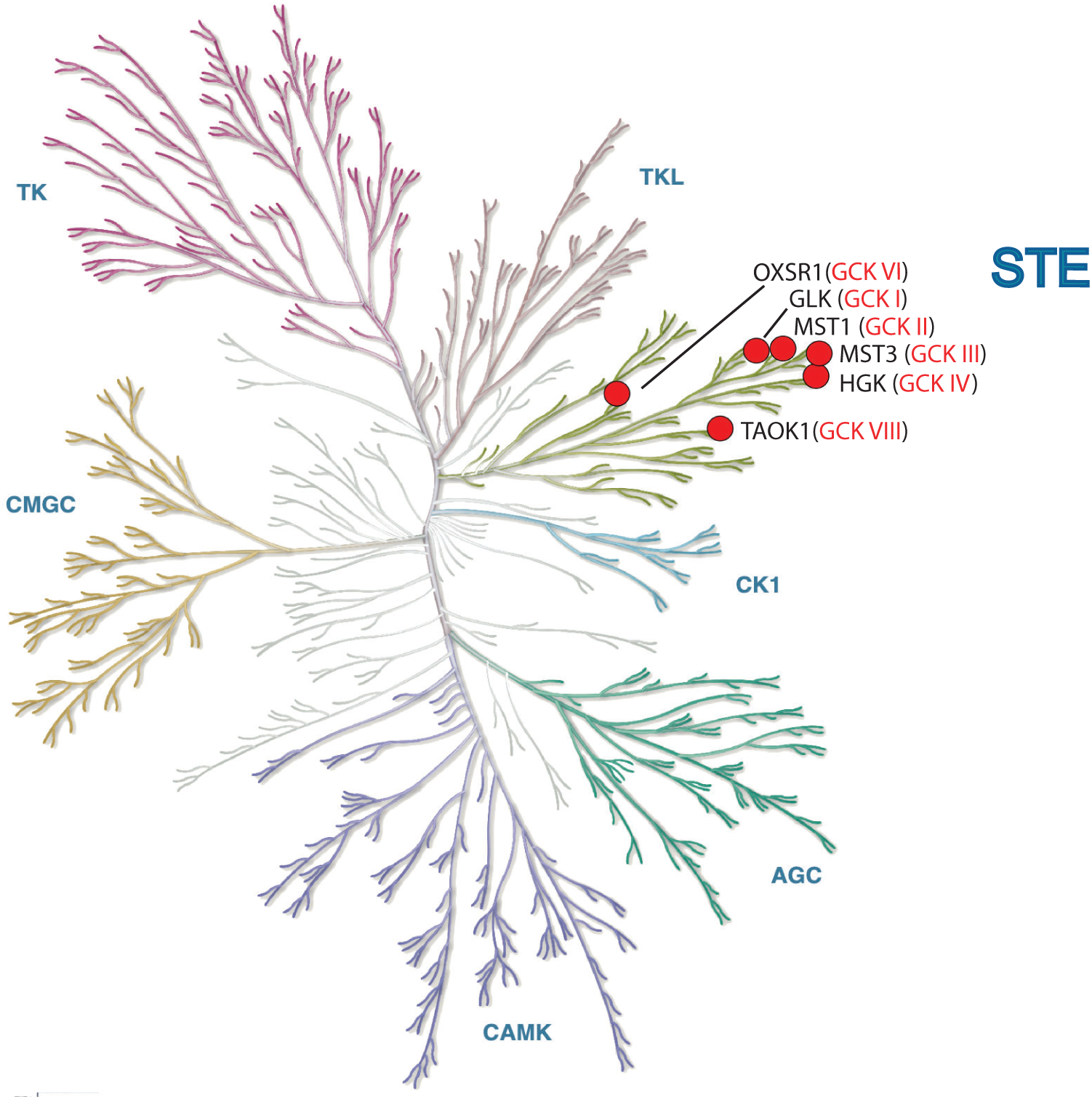
The GCK family kinases. Six GCK family members selected for PI3Kα activity screens. Their positions are highlighted on the evolutionary dendrogram of the human protein kinome.

**Fig. S2.**
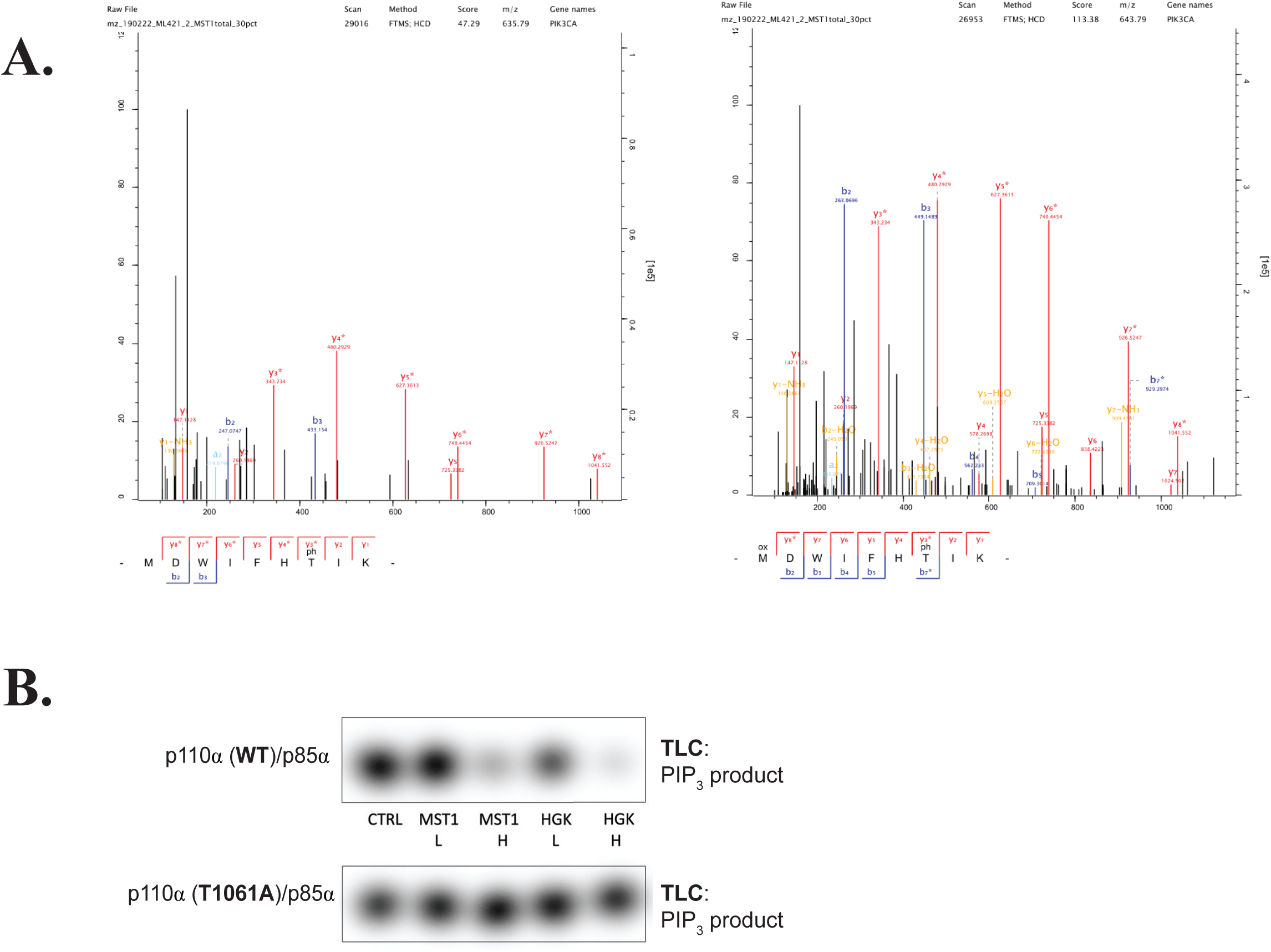
MST1/2 and HGK inhibit catalytic activity of p110*α* through phosphorylation at T1061. (A) Phosphopeptides containing T1061 were identified by MS/MS using purified p110α/p85α incubated with MST1 for 1h. Two versions of phosphopeptides were identified, with differences in methionine oxidation status. (B) Autoradiography of [^32^P]PIP_3_ production by the purified p110α (WT)/p85α or p110α (T1061A)/p85α after treatment with MST1 or HGK.

**Fig. S3.**
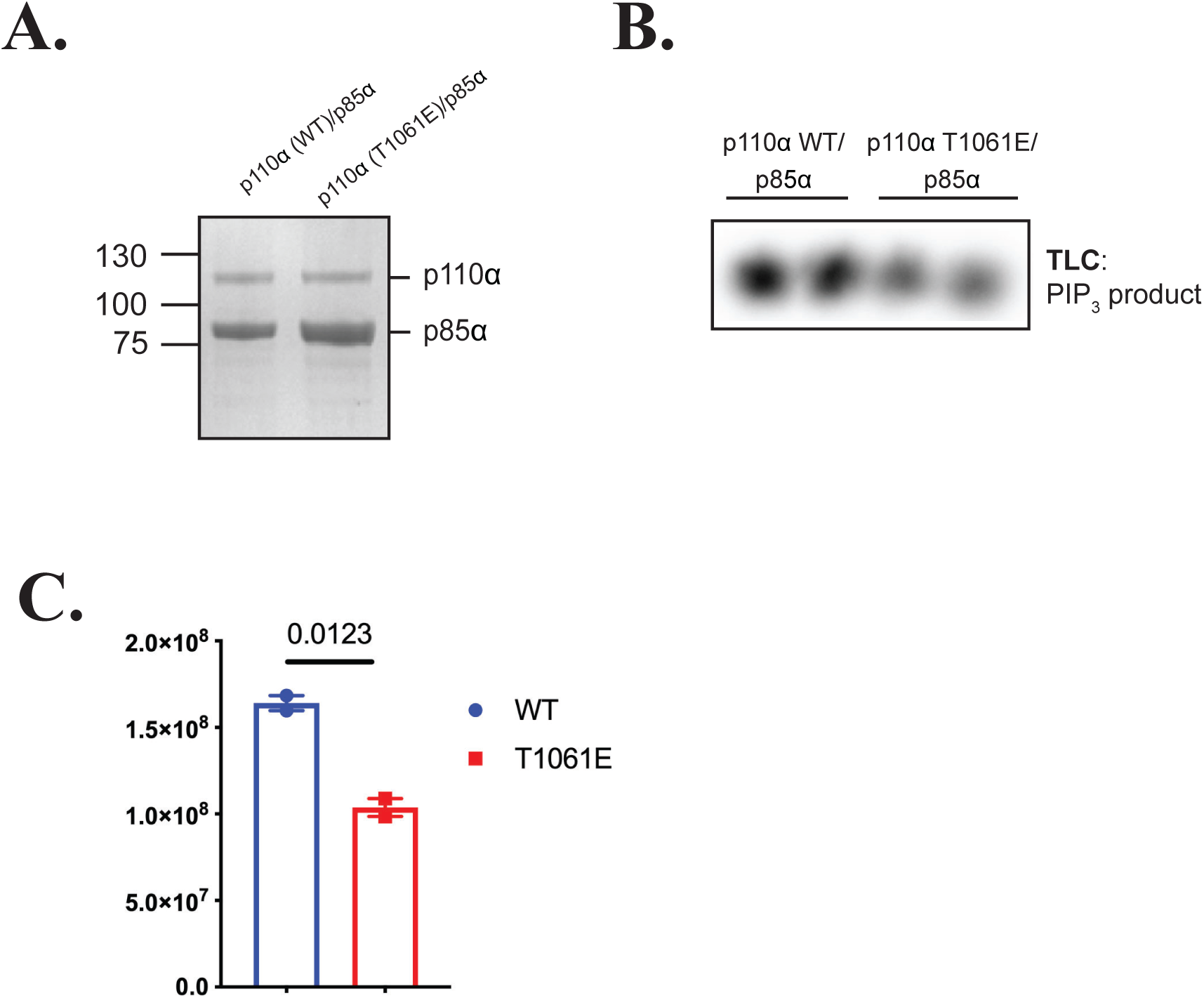
Characterization of the phosphomimetic T1061 p110a mutant. (A) Top: Coomassie stain comparison of purified recombinant p110a (WT)/His_6_-p85a and p110a (T1061E)/His_6_-p85a. (B) Second from top: Basal activities of p110α (WT) and the phosphomimetic mutant, p110α (T1061E) as measured by [^32^P]PIP_3_ production. (C) Bar graph of corresponding densitometries. Quantification of densitometries from (B). Data were represented as means ± SEMs. Significance, was calculated using student’s t test (N=2).

**Fig. S4.**
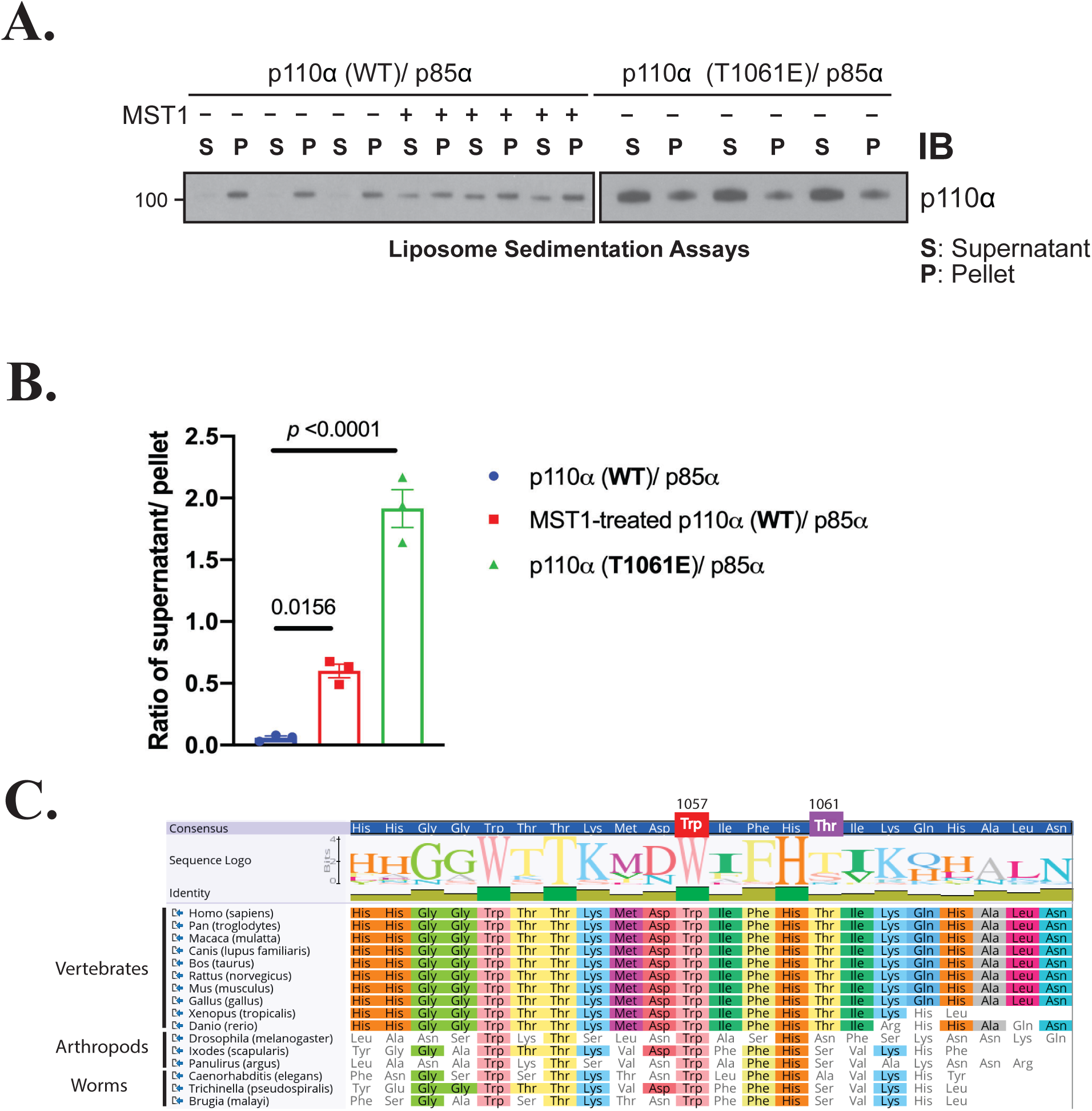
Phosphorylation of p110*α* by the Hippo kinases inhibits p110*α*’s interaction with membranes. (A) Liposome sedimentation assays of PI3Kα -/+ MST1 and untreated phosphomimetic mutant p110α (T1061E)/p85α. Immunoblots of p110α recovered from pellet (P) and supernatant (S). (B) Bar graph of corresponding densitometries from (A): as ratios of p110α recovery from supernatant over pellet. Data were represented as means ± SEMs. Significance was calculated using one-way ANOVA with Turkey’s multiple comparisons post-test (N=3). (C) Evolutionary alignment of the C-tails of p110α orthologs, highlighting W1057 and T1061.

**Fig. S5.**
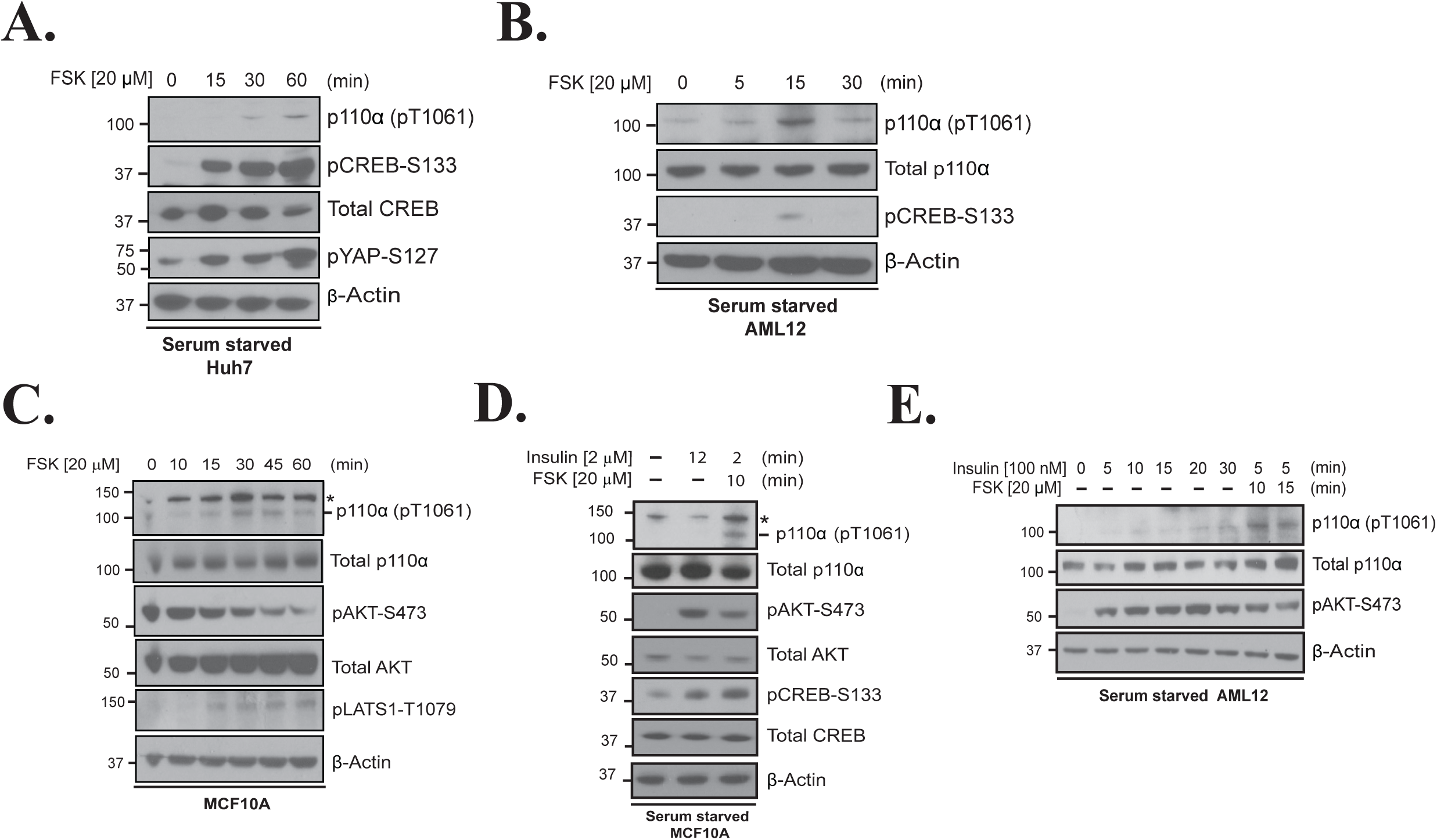
Activation of adenylyl cyclase promotes phosphorylation and inhibition of p110*α*. (A) Immunoblot for the indicated proteins using lysates from Huh7 cells that were serum starved for 12 h before stimulated with FSK [20 μM] for 0, 15, 30 and 60 min. (B) Immunoblot for the indicated proteins using lysates from AML12 cells that were serum starved for 12 h before stimulated with FSK [20 μM] for 0, 5, 15 and 30 min. (C) Immunoblot for the indicated proteins using lysates from MCF10A cells cultured in full medium that were stimulated with FSK [20 μM] for 0, 10, 15, 30, 45, and 60 min. (D) Immunoblot for the indicated proteins using lysates from MCF10A cells that were serum starved for 16 h, and then stimulated with DMSO, insulin [2 μM], or insulin [2 μM] followed by FSK [20 μM] for 10 min. *: non-specific band. (E) Immunoblot for the indicated proteins using lysates from AML12 cells that were serum starved for 12 h before stimulated with insulin [100 nM] for 0, 5, 10, 15, 20 and 30 min or insulin [100 nM] followed by FSK [20 μM] for 10 or 15 min.

**Fig. S6.**
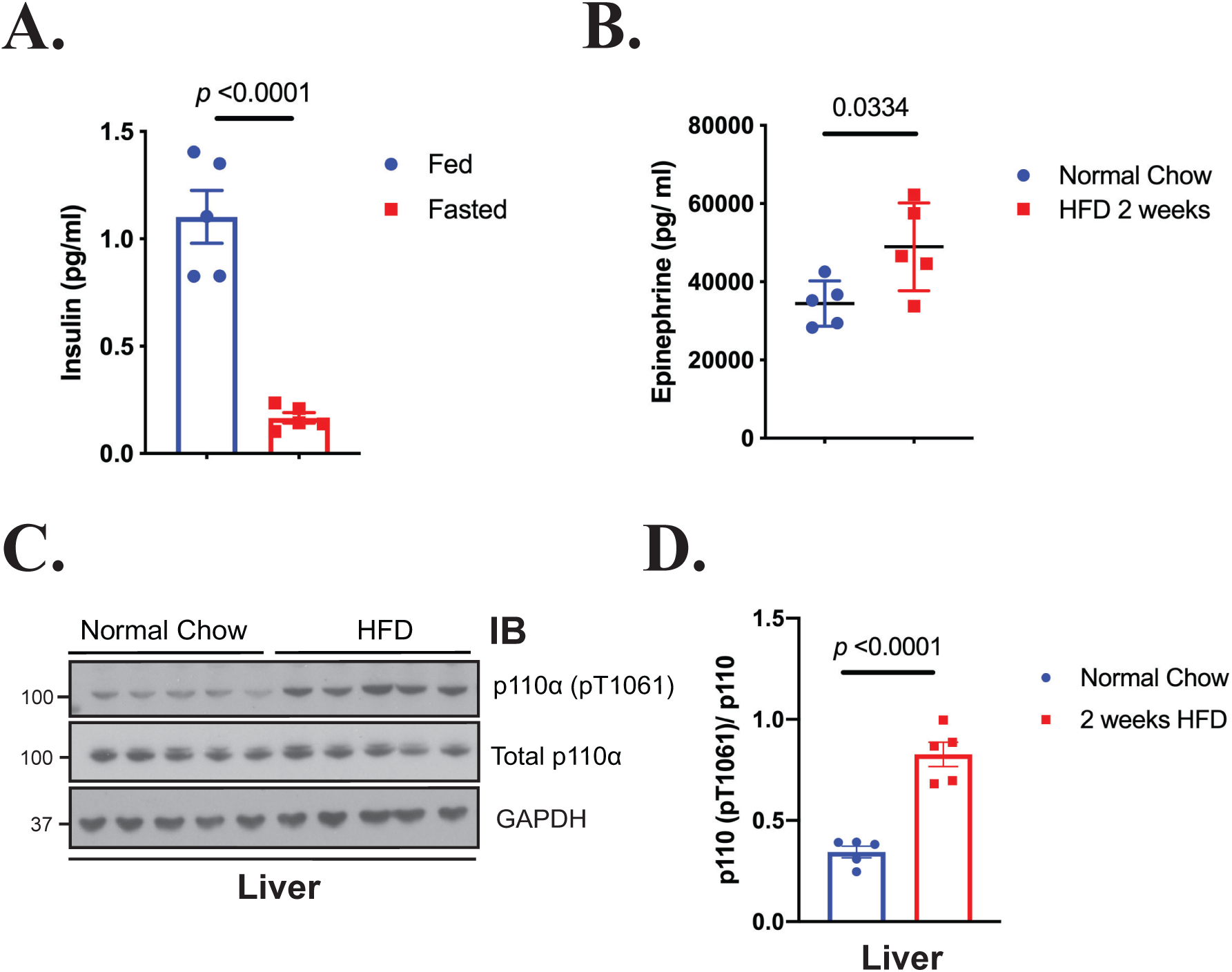
Epinephrine promotes phosphorylation of p110*α* in vivo. (A) Plasma insulin levels in WT mice that were fasted for 18 h (Fasted) and then euthanized or provided food for 4 h (Fed). Data were represented as means ± SEMs. Significance was calculated using student’s t test (N=5). (B) Plasma Epi levels in WT mice fed either normal chow or 60% high-fat diet (HFD) for 2 weeks. Data were represented as means ± SEMs. Significance was calculated using student’s t test (N=5). (C) Immunoblot for the indicated proteins using lysates from liver taken from WT mice fed normal chow or HFD for 2 weeks. (D) Quantification of the ratio of pT1061 to total p110α using band intensity from (C). Data is represented as means ± SEMs. Significance was calculated using student’s t test (N=5).

**Fig. S7.**
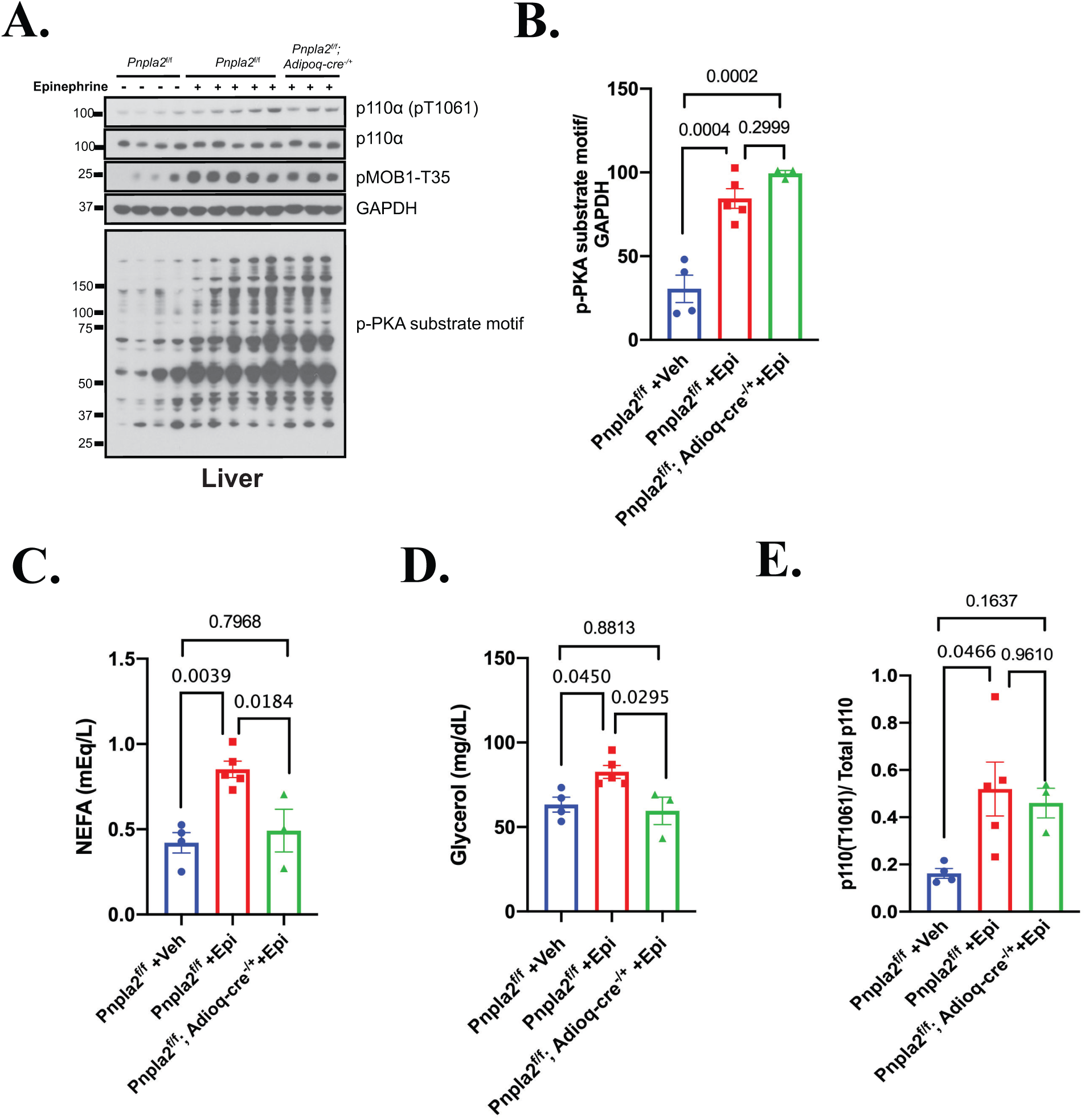
Effects of epinephrine on mice without adipocyte lipolysis. Lipolysis-deficient mice (Pnpla2^f/f^; Adipoq-Cre^+/-^) and littermate controls (Pnpla2^f/f^) were treated with vehicle (Veh, saline) or epinephrine (Epi) via I.P. injection (N=4, 5, 3). Blood and liver tissue were harvested 10 minutes later. (A) Markers of MST1/2 activation from liver lysates using Western blot. Quantification of combined PKA substrate motif lane normalized to GAPDH. Significance calculated using ANOVA with Turkey’s multiple comparisons post-test. Serum (C) non-esterified fatty acids (NEFA) and (D) glycerol from mice in A. Significance calculated using ANOVA with Turkey’s multiple comparisons post-test. (E) T1061 p110a phosphorylation normalized to total p110a. Significance calculated using ANOVA with Sidak’s multiple comparisons post-test. All data represented as means ± SEMs.

**Fig. S8.**
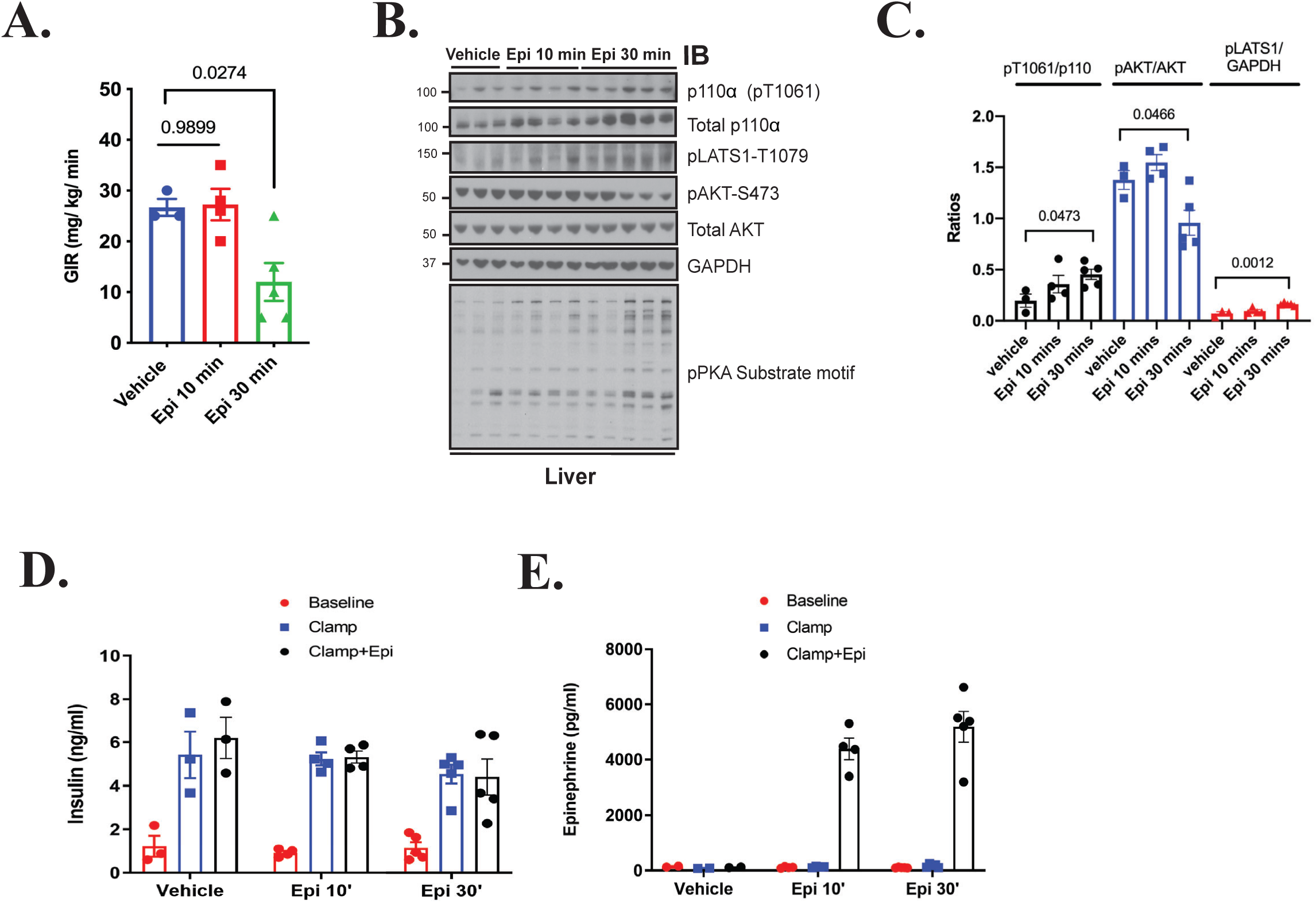
Epinephrine-mediated insulin resistance correlates with T1061 phosphorylation. (A) Steady-state glucose infusion rate (GIR) during a hyperinsulinemic-euglycemic clamp performed on WT rats. After 87 min, rats received an infusion of vehicle (normal saline, N=3) or Epi (0.75 μg/kg/min) for either 10 min (N=4) or 30 min (N=5). (B) Immunoblot for the indicated proteins using lysates from liver taken from WT rats in A. (C) Quantification of the ratio of pT1061 to total p110α, phosphorylated AKT to total AKT, and pLATS1 (T1079) to GAPDH using band intensity from (B). Significance calculated using ANOVA with Dunnett’s multiple comparisons to vehicle control. (D-E) Measurements of blood insulin and epinephrine levels during the hyperinsulinemic-euglycemic clamp. Data were represented as means ± SEMs.

**Fig. S9.**
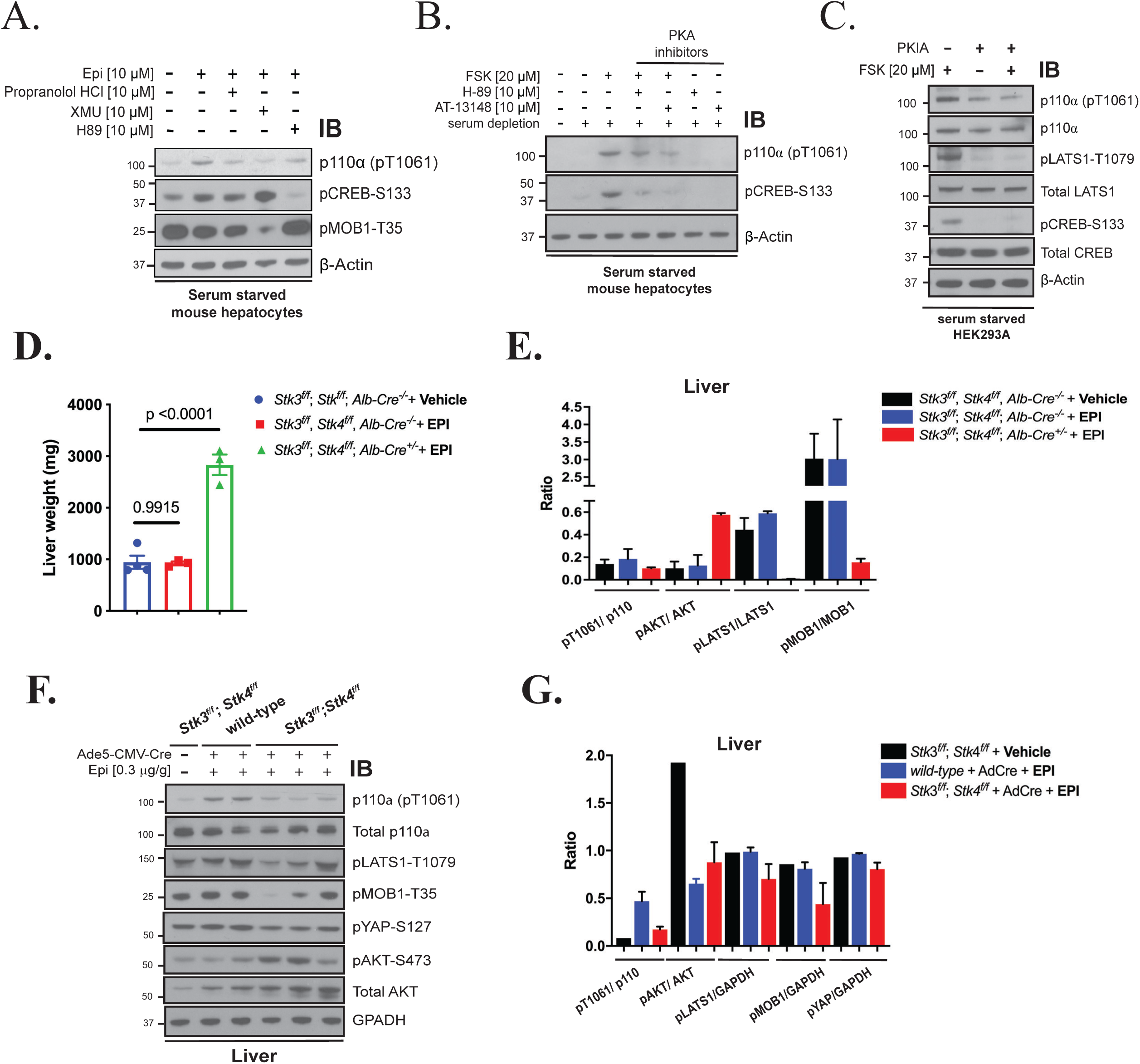
PKA and the Hippo kinases regulate FSK/ Epi-mediated p110*α* phosphorylation. (A) Immunoblot for the indicated proteins using lysates from serum starved primary mouse hepatocytes isolated from WT mice that were treated with Epi [10 μM], or Epi with Propranolol HCl [10 μM], XMU [10 μM], or H-89 [10 μM]. (B) Immunoblot for the indicated proteins using lysates from primary mouse hepatocytes that were serum starved 12 h before stimulated with DMSO, FSK [20 μM], FSK [20 μM] with two PKA inhibitors (H-89 and AT13148) [10 μM] or inhibitors alone. (C) Immunoblot for the indicated proteins using lysates from HEK293A cells transfected with control or PKIA vector f or 2 days before serum starved for 2 h, followed by stimulation with FSK [20 μM] for 15 min. (E) Quantification of the ratio of phosphorylated proteins over total proteins using band intensity from [Fig. 6E]. Data were represented as means ± SEMs. (F) Immunoblot for the indicated proteins using lysates from liver taken from Stk3^f/f^; Stk4^f/f^ or WT mice. Mice were injected with Ad5-CMV-Cre virus [10^9^] to induce liver-specific MST1/2 gene deletion, and then two weeks later, the mice were injected with vehicle (saline) or Epi [0.3ug/g] and liver were harvested.(G) Quantification of the ratio of phosphorylated proteins over total proteins using band intensity from (F).

**Fig. S10.**
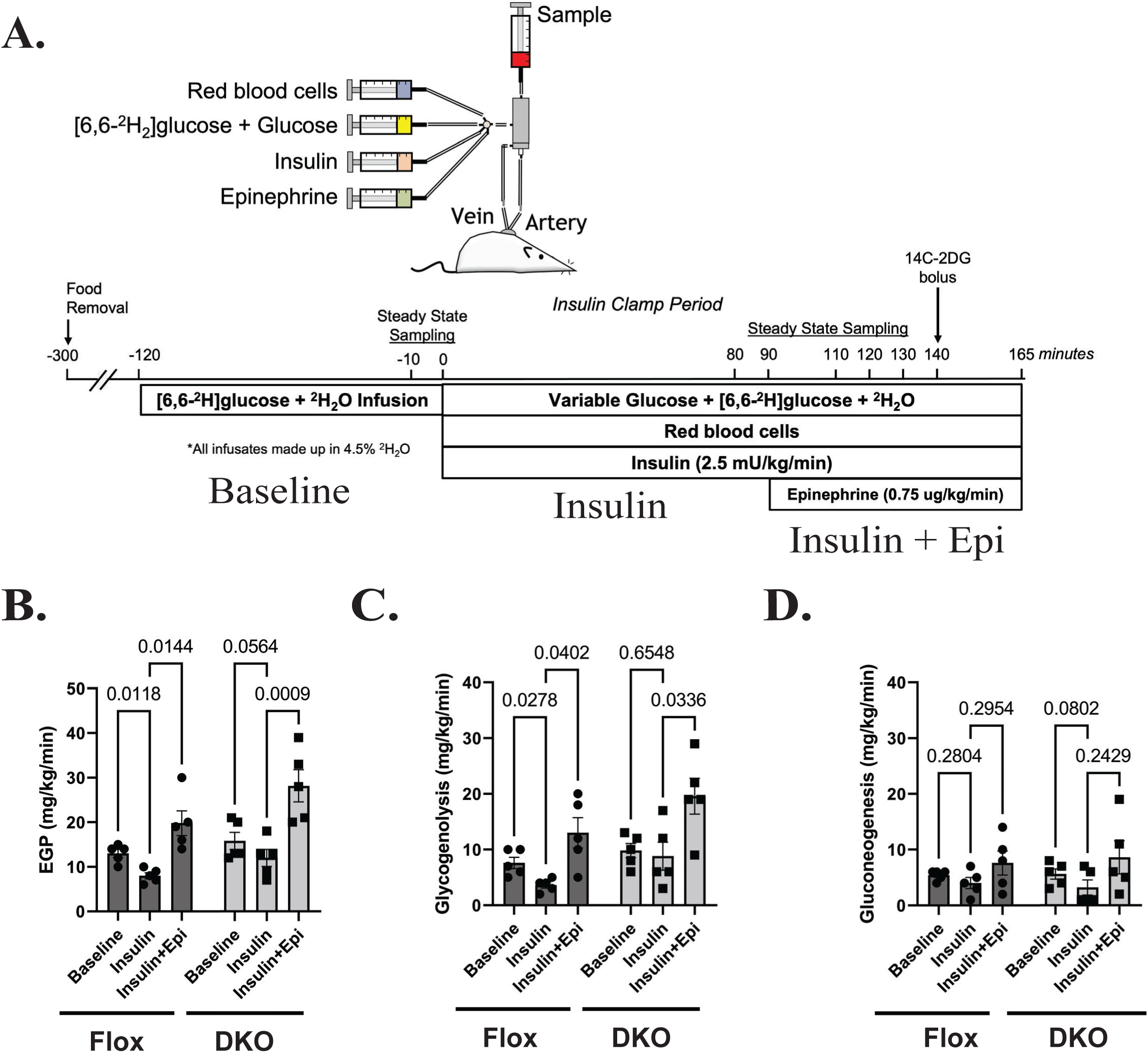
Epinephrine-mediated glycogenolysis is in tact in MST1/2 DKO mice under hyperinsulinemic-euglycemic conditions. (A) Schematic describing hyperinsulinemic-euglycemic-hyperepinephrinemic clamp protocol performed on Stk3^f/f^,Stk4^f/f^, Alb-Cre ^-/-^ (Flox, N=5) and Stk3^f/f^; Stk4^f/f^, Alb-Cre^+/-^ (DKO, N=5) mice. Rates of (B) endogenous glucose production (EGP), (C) glycogenolysis, and (D) gluconeogenesis during the clamp protocol. Comparisons made via Two-way ANOVA. Data presented as mean ± SEM.

